# Reversing Pdgfrβ Signaling Restores Metabolically Active Beige Adipocytes by Alleviating ILC2 Suppression in Aged and Obese Mice

**DOI:** 10.1101/2024.06.17.599436

**Authors:** Abigail M. Benvie, Daniel C. Berry

**Author notes:** Correspondence should be addressed to: Daniel Berry, Division of Nutritional Sciences, Cornell University, 526 Campus Rd., 307 Biotechnology Building, Ithaca, NY 14853, Contact.

## Abstract

**Objective:** Platelet Derived Growth Factor Receptor Beta (Pdgfrβ) suppresses the formation of cold temperature-induced beige adipocytes in aged mammals. We aimed to determine if deleting Pdgfrβ in aged mice could rejuvenate metabolically active beige adipocytes by activating group 2 innate lymphoid cells (ILC2), and whether this effect could counteract diet-induced obesity-associated beige fat decline.

**Methods:** We employed Pdgfrβ gain-of-function and loss-of-function mouse models targeting beige adipocyte progenitor cells (APCs). Our approach included cold exposure, metabolic cage analysis, and age and diet-induced obesity models to examine beige fat development and metabolic function under varied Pdgfrβ activity.

**Results:** Acute cold exposure alone enhanced metabolic benefits in aged mice, irrespective of beige fat generation. However, Pdgfrβ deletion in aged mice reestablished the formation of metabolically functional beige adipocytes, enhancing metabolism. Conversely, constitutive Pdgfrβ activation in young mice stymied beige fat development. Mechanistically, Pdgfrβ deletion upregulated IL-33, promoting ILC2 recruitment and activation, whereas Pdgfrβ activation reduced IL-33 levels and suppressed ILC2 activity. Notably, diet-induced obesity markedly increased Pdgfrβ expression and Stat1 signaling, which inhibited IL-33 induction and ILC2 activation. Genetic deletion of Pdgfrβ restored beige fat formation in obese mice, improving whole-body metabolism.

**Conclusion:** This study reveals that cold temperature exposure alone can trigger metabolic activation in aged mammals. However, reversing Pdgfrβ signaling in aged and obese mice not only restores beige fat formation but also renews metabolic function and enhances the immunological environment of white adipose tissue (WAT). These findings highlight Pdgfrβ as a crucial target for therapeutic strategies aimed at combating age- and obesity-related metabolic decline.

## INTRODUCTION

Aging is associated with the development of obesity, increasing the risk of developing type 2 diabetes and cardiometabolic diseases[1; 2]. Obesity—excess fat mass—is often associated with altered energy balance and the consumption of nutrient-low calories, which drives white adipose tissue dysfunction. This dysfunction prevents glucose uptake, reduces adipocyte lipid storage, enables ectopic fat deposition, and augments blood lipid levels[1; 3]. Accordingly, identifying molecular regulators opposing fat mass accumulation could enable metabolic changes in aged and obese individuals.

Cold temperature exposure reduces fat mass and improves systemic metabolism and energy expenditure[4; 5]. It does so, in part, by recruiting and activating thermogenic beige adipocytes within white adipose tissues (WAT) to burn free fatty acids and glucose. Like brown adipocytes, beige fat cells are multilocular and have specialized mitochondria that express uncoupling protein 1 (Ucp1) to collapse the proton gradient to release heat[6]. Indeed, stimulated beige adipocytes have been identified in cold-exposed (15°C) adult humans[7–11]. Markedly, the capacity to generate beige adipocytes reduces the risk of developing obesity and metabolic disease[12]. For instance, beige fat transplantation studies have shown host recipient changes in glucose metabolism, insulin sensitivity, reduced adiposity, and lower body mass[13; 14]. Moreover, the ability to recruit or activate beige adipose tissue is associated with changes in body fat distribution, such as reducing visceral fat[4]. On the other hand, preventing thermogenic fat formation increases the risk of developing metabolic diseases[15], suggesting that thermogenic fat has metabolic utility and underscoring the ability to generate more. Indeed, recent efforts to engage human thermogenic fat activation have focused on modulating ambient temperature to reduce body weight and attenuate metabolic pathologies.

To generate beige adipocytes, cold temperatures trigger the adipogenic action of WAT-resident perivascular smooth muscle cells. These beige fat progenitors resemble mural cells and express key vascular markers, such as smooth muscle actin (Sma/Acta2), Sm22 (Tagln), Myh11, and CD81[16–18]. However, the ability of cold temperatures to generate mammalian beige adipocytes declines with age, dampening the therapeutic promise of beige fat within aging populations[19–23]. Moreover, patients with obesity also show a decline in cold-induced beige fat development[24]. Remarkably, the decline in beige fat development can be observed in humans starting in their mid-30s, before the decline of age-related tissue homeostasis and obesity[23]. Cold-induced beige adipocyte failure can be attributed to several pathways and processes, such as cellular senescence, WAT fibrosis, inflammation, and mitophagy[20; 21; 25-27]. Our previous study identified the age-dependent upregulation of Pdgfrβ within senescent beige progenitor cells. Deleting Pdgfrβ in aged mice restores Ucp1+ multilocular beige adipocytes within WATs. Conversely, beige adipocyte development was diminished in young mice harboring a constitutively activated Pdgfrβ, mirroring an aging phenotype. Changes in Pdgfrβ expression were associated with lower Th2-cytokine signaling pathways[22]. Yet, several limitations existed, such as whether beige adipocytes induced by Pdgfrβ deletion are metabolically active and how Pdgfrβ remodels the immunological niche. Moreover, does Pdgfrβ activation contribute to obesity-associated beige adipocyte decline?

A growing body of evidence suggests the metabolic benefits of repeated cold exposure in humans, such as boosting energy expenditure and promoting whole-body fatty acid and glucose use[10; 28-30]. Here, we use Pdgfrβ signaling to identify the contribution of metabolically active beige fat development to energy regulation and metabolism in aging and obese mice. This study shows that aging and obesity reprogram beige adipocyte progenitors by upregulating Pdgfrβ signaling to suppress the ILC2 recruitment preventing beige fat development and reducing energy expenditure.

## RESULTS

### Cold exposure can improve age-associated changes in physiology

To evaluate the impact of aging and beige fat decline on cold-induced metabolism, we exposed 2-month-(young) and 12-month-old (aged) male and female mice to room temperature (RT) (23°C) or cold exposure (CE) (6.5°C) for seven days (Figure 1A). We first examined whether CE could alter several physiological parameters in the aged setting. As expected, RT-aged male and female mice were heavier than RT-juvenile mice (Figure 1B, C). Aging in male mice at RT was associated with impaired glucose clearance compared to their young RT counterparts, whereas aged RT female mice appeared protected against age-related changes in glucose clearance (Figure 1D-G). However, in response to CE, we observed a significant reduction in body weight in both young and aged male and female mice (Figure 1B, C). Additionally, CE improved glucose clearance in aged male and female mice, indicating a cold-induced metabolic benefit independent of beige fat formation (Figure 1D-G). Critically, CE significantly reduced serum triglycerides and cholesterol levels regardless of age (Figure 1H-K).

**Figure 1:**
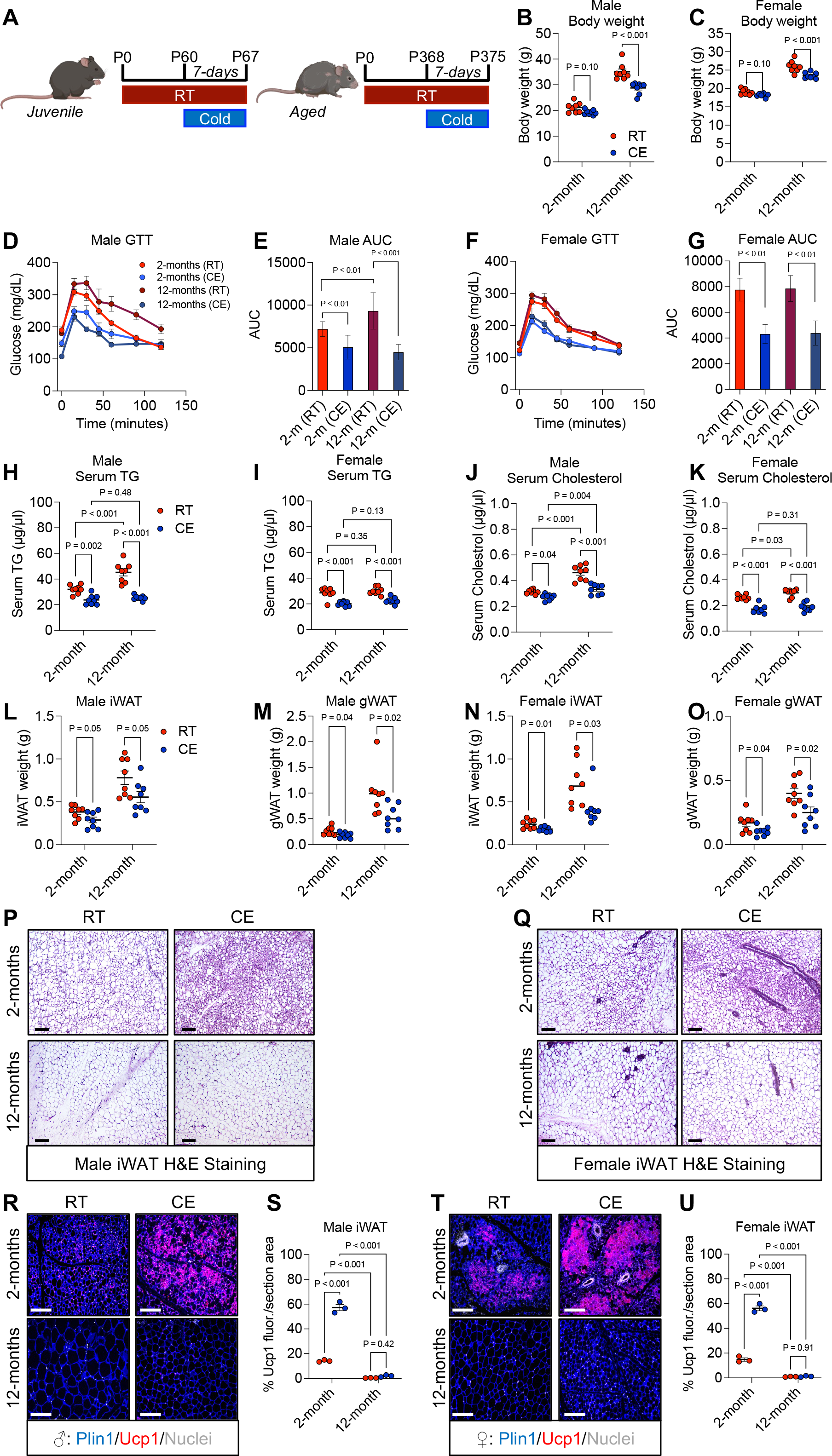
Cold exposure improves metabolic parameters in aged mice, independent of beige fat formation. **A.** Experimental timeline to evaluate RT and CE on aging mice **B, C.** Body weight of male (B) and female (C) mice maintained at RT or CE exposed at 2 months (young) or 12 months (aged) (n = 8 biologically independent mice). **D, E.** Intraperitoneal glucose tolerance test performed on young or aged male mice at RT or after seven days of CE (n = 8 biologically independent mice). **F, G.** Intraperitoneal glucose tolerance test performed on young or aged female mice at RT or after seven days of CE (n = 8 biologically independent mice). **H, I.** Serum triglyceride (TG) levels in male (H) and female (I) mice maintained at RT or CE at 2 or 12 months (n = 8 biologically independent mice). **J, K.** Serum cholesterol levels in male (J) and female (K) mice maintained at RT or CE at 2 or 12 months (n = 8 biologically independent mice). **L, M.** Inguinal (iWAT) and pergonadal (gWAT) tissue weight from male mice maintained at RT or CE at 2 or 12 months of age (n = 8 biologically independent mice). **N, O.** iWAT and gWAT tissue weight from female mice maintained at RT or CE at 2 or 12 months of age (n = 8 biologically independent mice). **P.** Representative images of H&E staining of iWAT sections from male mice maintained at RT or CE at 2 or 12 months of age **Q.** Representative images of H&E staining of iWAT sections from female mice maintained at RT or CE at 2 or 12 months of age. **R, S.** Representative images of Ucp1 immunostaining (**R**) and quantification (**S**) of iWAT sections from male mice maintained at RT or CE at 2 or 12 months of age (n = 3 biologically independent mice). **T, U.** Representative images of Ucp1 immunostaining (**R**) and quantification (**S**) of iWAT sections from female mice maintained at RT or CE at 2 or 12 months of age (n = biologically independent mice). Data are means with individual data points ±S.E.M. Statistical significance was determined using unpaired two-tailed Student’s *t* test (B, C,L, M, N and O) and one-way ANOVA (E, G, H-K, S, and U). Scale bar = 100 µm.

Next, we evaluated whether CE could reduce adiposity in aged mice. As expected, aged male and female mice had expanded inguinal (iWAT) and perigonadal (gWAT) adipose tissues compared to young mice (Figure 1L-O). Nevertheless, CE reduced adipose depot weight in both young and aged male and female mice (Figure 1L-O). Consistent with this observation, histological staining revealed that CE decreased the size of unilocular white adipocytes in iWAT and gWAT depots compared to RT-aged controls (Figure 1P-Q and Supplementary Fig. 1A, B). To confirm these findings, we measured adiponectin, an adipokine associated with WAT size and health[31]. We observed downregulation of serum adiponectin levels in RT-aged male and female mice, presumably due to increased WAT size and age-related decline in tissue function (Supplementary Fig. 1C, D). Conversely, CE boosted adiponectin levels in both young and aged male and female mice, suggesting that CE improves adipose tissue function and systemic metabolism, independent of beige fat biogenesis (Supplementary Fig. 1C, D).

In response to CE, young male and female mice readily developed Ucp1+ beige adipocytes within the dorsal lumbar region of iWAT depots (Figure 1P-U). Notably, only female mice developed pronounced CE-induced Ucp1+ beige fat formation in perigonadal white adipose tissue (gWAT) depots (Supplementary Fig. 1E-H). Moreover, the appearance of beige adipocytes in young mice was associated with the induction of a thermogenic program (Supplementary Fig. 1I). In contrast, aged male and female mice rarely developed beige adipocytes in either iWAT or gWAT depots (Figure 1P-U and Supplementary Fig. 1E-H). Aging significantly diminished the induction of Ucp1 and other thermogenic genes within iWAT depots, underscoring an age-associated decline in beige fat biogenesis (Supplementary Fig. 1I). Overall, our results demonstrate an age-associated decline in beige fat biogenesis in both male and female mice.

### Pdgfrβ deletion restores metabolically functional beige fat in aged mice

We previously identified the age-associated upregulation of Pdgfrβ mRNA and protein expression within WAT[22]. Correspondingly, deleting Pdgfrβ in aged APCs restored beige fat appearance. To extend these findings, we sought to understand whether Pdgfrβ-KO induced metabolically functional beige adipocytes and, if so, whether these beige adipocytes bolster metabolism in aged mice. To test this notion, we combined the Pdgfrβ^fl/fl^ conditional mouse model with the tamoxifen (TMX)-inducible Sma-Cre^ERT2^[22; 32]. Aged mice (12-months-old) were administered TMX to delete Pdgfrβ within Sma+ beige precursors[16]. After TMX induction, we subsequently CE mice for seven days and evaluated metabolic parameters (Figure 2A). As expected, CE-aged Pdgfrβ-KO male and female mice upheld homothermic responses compared to age-matched controls (Figure 2B, C). Strikingly, aged Pdgfrβ-KO male and female mice displayed a significant increase in energy expenditure, indicating that Pdgfrβ deletion generated functional beige adipocytes (Figure 2D, E and Supplementary Fig. 2A, B). In addition to elevated energy expenditure, Pdgfrβ-KO mice had a lower respiratory exchange ratio (RER), suggesting a macronutrient shift to lipids (Figure 2F, G). However, changes in RER appeared to be independent of food intake (Figure 2H, I). Next, we assessed if Pdgfrβ deletion improved blood glucose clearance after CE by performing an intraperitoneal glucose tolerance test. After CE, the blood glucose peak and decline were significantly enhanced in Pdgfrβ-KO male and female mice (Figure 2J, K). Consistently, we observed improvements in other serum indicators, such as triglycerides and cholesterol levels in cold-exposed Pdgfrβ-KO mice (Supplementary Fig. 2C-F). Together, deleting Pdgfrβ in SMA cells appears to shift several physiological surrogates of beige fat development and may enhance CE-induced metabolic effects.

**Figure 2:**
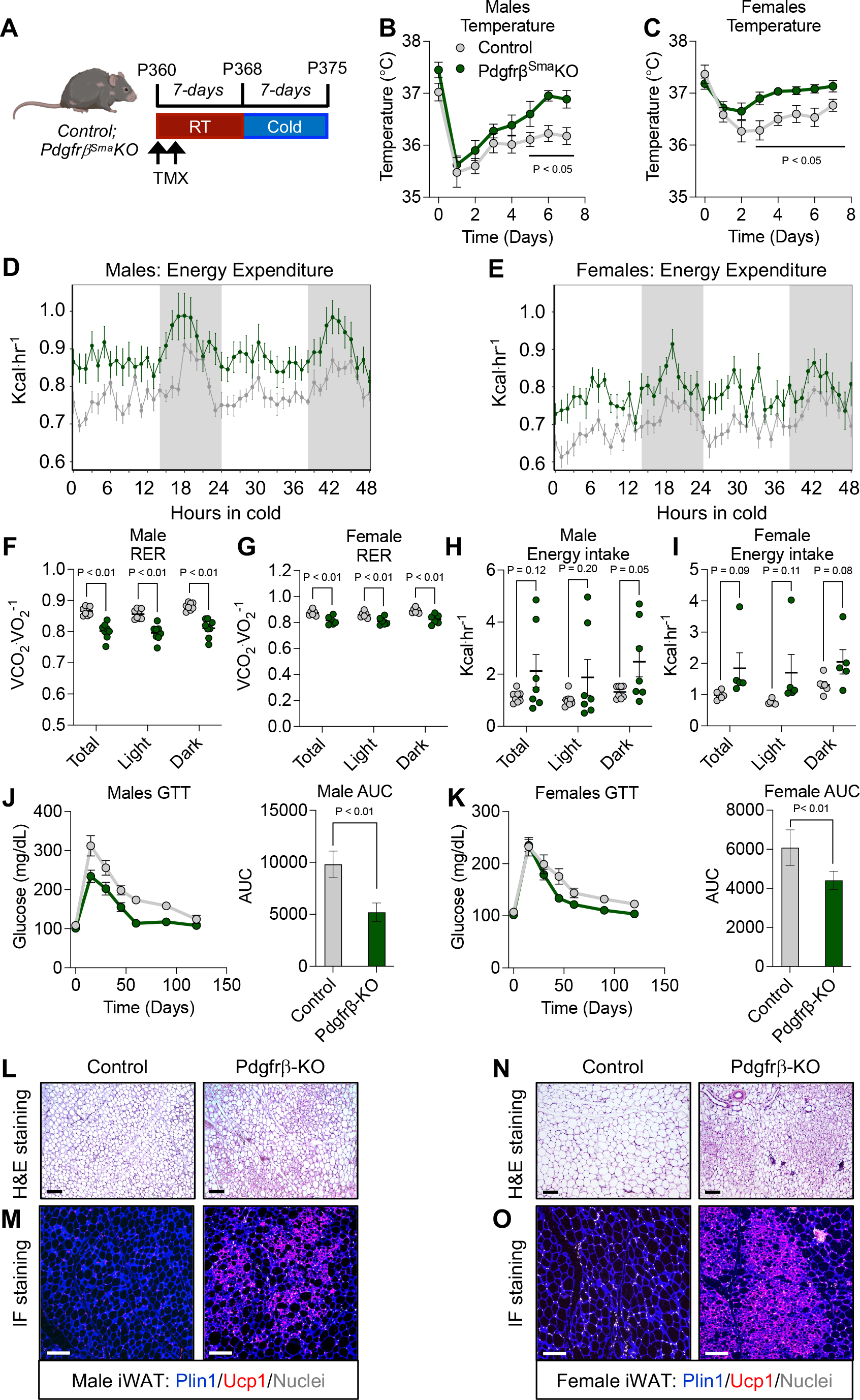
Deletion of Pdgfrβ improves cold temperature induced metabolic parameters and beige fat in aged mice. **A.** Experimental design to evaluate Control and Pdgfrβ-KO aging mice. **B, C.** Rectal temperature of male (B) and female (C) Control and Pdgfrβ-KO aged mice throughout CE (n = 8 biologically independent mice). **D, E.** Energy expenditure of Control and Pdgfrβ-KO aged male (D) and female (E) mice acclimated to CE (n = 6 biologically independent mice). **F, G.** Respiratory exchange ratio (RER) of Control and Pdgfrβ-KO aged male (F) and female (G) mice acclimated to CE (n = 6-8 biologically independent mice). **H, I.** Energy intake of Control and Pdgfrβ-KO aged male (H) and female (I) mice acclimated to CE (n = 6-8 biologically independent mice). **J.** Intraperitoneal glucose tolerance test performed on Control and Pdgfrβ-KO aged male mice after seven days of CE. Right: Area under the curve (AUC) was calculated (n = 8 biologically independent mice). **K.** Intraperitoneal glucose tolerance test performed on Control and Pdgfrβ-KO aged female mice after seven days of CE (n = 8 biologically independent mice). Right: Area under the curve (AUC) was calculated. **L.** Representative images of H&E staining of iWAT sections from Control and Pdgfrβ-KO male mice maintained at CE for seven days. **M.** Representative images of Ucp1 immunostaining of iWAT sections from Control and Pdgfrβ-KO male mice maintained at CE for seven days. **N.** Representative images of H&E staining of iWAT sections from Control and Pdgfrβ-KO female mice maintained at CE for seven days. **O.** Representative images of Ucp1 immunostaining of iWAT sections from Control and Pdgfrβ-KO female mice maintained at CE for seven days. Data are means with individual data points ±S.E.M. Statistical significance was determined using unpaired two-tailed Student’s *t* test (B, C, and F-K). Scale bar = 100 µm.

Even though we observed changes in energy expenditure, body weight remained relatively constant in male control and mutant mice (Supplementary Fig. 2G). However, we observed that Pdgfrβ deletion slightly reduced body weight in female mice (Supplementary Fig. 2H). Cold temperature exposure was also associated with reduced adiposity in mutant males and females compared to controls (Supplementary Fig. 2I-L). Yet, we did not observe significant differences in non-adipose tissue organ weight (Supplementary Fig. 2M, N). Likewise, we did not observe changes in skeletal muscle shivering genes[33] (Supplementary Fig. 2O). Hematoxylin and eosin staining and Ucp1 immunostaining of aged control dorsal lumbar iWAT[34] sections after CE revealed minimal Ucp1+ multilocular beige adipocytes. In contrast, Pdgfrβ deletion significantly restored iWAT and gWAT beige fat development in aged male and female mice (Figure 2L-O and Supplementary Fig. 2P-U). Moreover, gene expression revealed that deleting Pdgfrβ restored thermogenic gene expression in iWAT depots after CE in aged mice (Supplementary Fig. 2V, W). Overall, deleting Pdgfrβ in aged mice restores functional beige adipocytes to improve metabolism.

### Pdgfrβ activation blunts the development of metabolically active beige fat in young mice

The data suggest that aging prevents beige fat generation, but Pdgfrβ deletion can restore beige fat development and boost several health benefits of CE. To realize the contribution of Pdgfrβ-induced blockade on beige fat development and metabolism, we impaired beige adipocyte formation, independent of brown fat, in young two-month-old mice to mimic aging. To do so, we previously generated a mouse model in which a constitutively Pdgfrβ activated allele is expressed within SMA+ beige fat progenitors[22]. In this model, the Pdgfrβ^D849V^ conditional mouse model is combined with the Sma-Cre^ERT2^ driver to create Pdgfrβ^SmaD849V^. Notably, this Cre-inducible constitutively active Pdgfrβ allele is controlled by the Pdgfrβ promoter and expressed at endogenous levels, but basal Pdgfrβ signaling is elevated[35–37]. Because we observed Pdgfrβ acting in an age-dependent manner to regulate beige fat formation negatively, we evaluated the necessity of Pdgfrβ in young mice using the Pdgfrβ-KO mouse model as a control. To induce recombination in young mice, we administered one dose of TMX for two consecutive days to delete Pdgfrβ or activate the Pdgfrβ^D849V^ allele. After the TMX washout[38], we cold exposed male and female mice for seven days (Figure 3A). As anticipated, juvenile control and Pdgfrβ-KO mice maintained similar body temperature defenses in response to CE. However, young mice harboring the Pdgfrβ^D849V^ could not defend their body temperature during CE (Figure 3B, C). In agreement with impaired homothermic control, after three days of CE, Pdgfrβ^D849V^ male and female mice had significantly lower energy expenditure than control and Pdgfrβ-KO male and female mice (Figure 3D, E and Supplementary Fig. 3A, B). We also observed a higher respiratory exchange ratio (RER), suggesting a macronutrient switch to primarily carbohydrate utilization (Figure 3F, G). However, changes in Pdgfrβ^D849V^ RER appeared to be independent of food intake (Figure 3H, I). Since brown adipose tissue is the primary site of thermogenesis and energy regulation[39], we have previously evaluated BAT characteristics in these mice and found no significant differences in thermogenic genes or histological appearance[22]. Moreover, Sma+ cells do not serve as a brown adipocyte precursor[15; 22].

**Figure 3:**
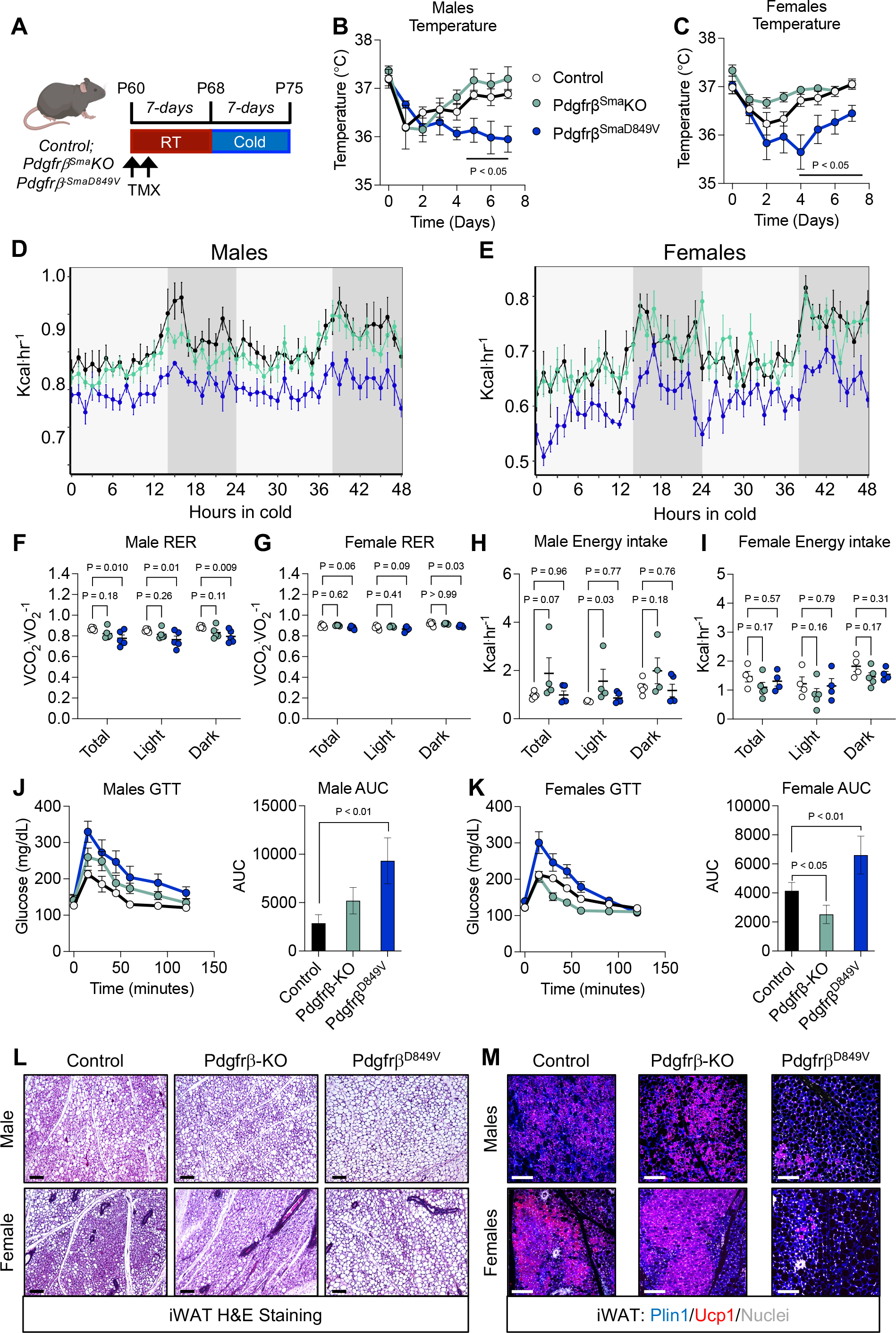
Constitutive activation of Pdgfrβ diminishes cold temperature induced metabolic parameters and blocks beige fat development in young mice. **A.** Experimental design to evaluate Control, Pdgfrβ-KO, and PdgfrβD849V young (2-month-old) mice after CE. **B, C.** Rectal temperature of male (B) and female (C) Control, Pdgfrβ-KO, and PdgfrβD849V young mice throughout CE (n = 6 biologically independent mice). **D, E.** Energy expenditure of Control, Pdgfrβ-KO, and PdgfrβD849V young male (D) and female (E) mice acclimated to CE (n = 5 biologically independent mice). **F, G.** Respiratory exchange ratio (RER) of Control, Pdgfrβ-KO, and PdgfrβD849V young male (F) and female (G) mice acclimated to CE (n = 5 biologically independent mice). **H, I.** Energy intake of Control, Pdgfrβ-KO, and PdgfrβD849V young male (H) and female (I) mice acclimated to CE (n = 5 biologically independent mice). **J.** Intraperitoneal glucose tolerance test performed on Control, Pdgfrβ-KO, and PdgfrβD849V young male mice after seven days of CE. Right: Area under the curve (AUC) was calculated (n = 6 biologically independent mice). **K.** Intraperitoneal glucose tolerance test performed on Control, Pdgfrβ-KO, and PdgfrβD849V young female mice after seven days of CE (n = 6 biologically independent mice). Right: Area under the curve (AUC) was calculated. **L.** Representative images of H&E staining of iWAT sections from Control, Pdgfrβ-KO, and PdgfrβD849V young male (top) and female (bottom) mice maintained at CE for seven days. **M.** Representative images of Ucp1 immunostaining of iWAT sections from Control, Pdgfrβ-KO, and PdgfrβD849V young male (top) and female (bottom) mice maintained at CE for seven days. Data are means with individual data points ±S.E.M. Statistical significance was determined using unpaired two-tailed Student’s *t* test (B, C, and F-K). Scale bar = 100 µm.

Next, we evaluated whether reduced energy expenditure was associated with changes in glucose responses. Consistent with this notion, the blood glucose peak and subsequent decline in Pdgfrβ^D849V^ male and female mice remained elevated throughout a glucose tolerance test (Figure 3J, K). Pdgfrβ^D849V^ dampened glucose clearance was also associated with elevated serum triglyceride and cholesterol levels (Supplementary Fig. 3C-F). While body weight remained comparable between control and mutant female mice, male mice harboring the Pdgfrβ^D849V^ mutation were slightly heavier (Supplementary Fig. 3G, H). In addition, adiposity remained higher in Pdgfrβ^D849V^ male and female mice compared to control and Pdgfrβ-KO mice (Supplementary Fig. 3I-L). However, non-adipose tissue organ weight remained comparable between control, Pdgfrβ-KO, and Pdgfrβ^D849V^ (Supplementary Fig. 3M, N). As previously observed[22], histological analysis and Ucp1 immunostaining revealed a paucity of beige fat development throughout Pdgfrβ^D849V^ iWAT depots (Figure 3L, M). Furthermore, Ucp1-positive beige adipocyte development within female gWAT depots was significantly blunted (Supplementary Fig. 3O-V). Thus, activating Pdgfrβ prevents functional beige fat formation in young mice to blunt metabolism.

### Pdgfrβ signaling coordinates ILC2 accrual

We previously found that Pdgfrβ expression negatively correlated with IL-33 expression to suppress Th2 cytokine signaling[22]. However, the regulation of IL-33 by Pdgfrβ signaling and its role in ILC2 accrual required further investigation. To determine if Pdgfrβ influences IL-33 bioavailability, we assessed IL-33 mRNA expression and secretion from beige APCs isolated from iWAT depots of aged control and mutant mice. Indeed, IL-33 expression was elevated within iWAT depots of Pdgfrβ-KO mice (Figure 4A). Strikingly, IL-33 secretion was nearly undetectable in aged control mice, but Pdgfrβ ablation significantly elevated IL-33 protein levels within the media (Figure 4B).

**Figure 4:**
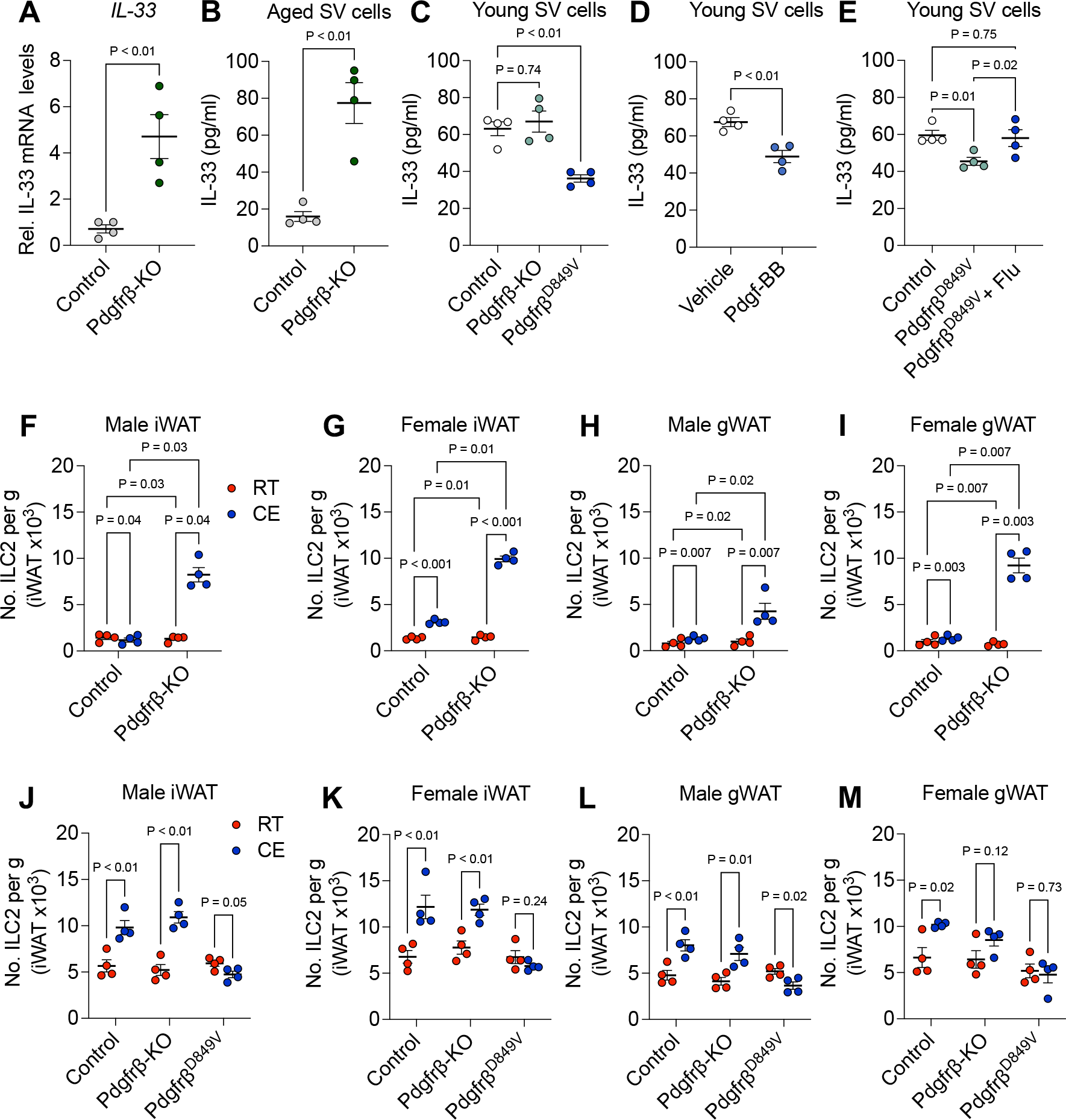
Pdgfrβ associates with IL-33 bioavailability and prevents ILC2 accrual. **A.** Relative mRNA expression levels of IL-33 expression within iWAT depots from aged Control and Pdgfrβ-KO male mice maintained at RT (n = 4 biologically independent mice). **B.** IL-33 protein levels (ELISA) in cell culture media from iWAT SV cells isolated from aged Control and Pdgfrβ-KO male mice maintained at RT (n = 4 biologically independent mice). **C.** IL-33 protein levels (ELISA) in cell culture media from iWAT SV cells isolated from young Control, Pdgfrβ-KO, and PdgfrβD849V male mice maintained at RT (n = 4 biologically independent mice). **D.** IL-33 protein levels (ELISA) in cell culture media from iWAT SV cells isolated from young Control, maintained at RT treated with vehicle or Pdgf-BB (50 ng/ml) for 48 hours (n = 4 biologically independent mice). **E.** IL-33 protein levels (ELISA) in cell culture media from iWAT SV cells isolated from young Control and PdgfrβD849V male mice maintained at RT treated with vehicle or fludarabine (1µM) for 48 hours (n = 4 biologically independent mice). **F, G.** FACS quantification of ILC2 abundance in iWAT depots from aged Control and Pdgfrβ-KO male (F) and female (G) mice after CE (n = 4 biologically independent mice). **H, I.** FACS quantification of ILC2 abundance in gWAT depots from aged Control and Pdgfrβ-KO male (H) and female (I) mice after CE (n = 4 biologically independent mice). **J, K.** FACS quantification of ILC2 abundance in iWAT depots from young Control, Pdgfrβ-KO, and PdgfrβD849V male (J) and female (K) mice after CE (n = 4 biologically independent mice). **L, M.** FACS quantification of ILC2 abundance in gWAT depots from young Control, Pdgfrβ-KO, and PdgfrβD849V male (L) and female (M) mice after CE (n = 4 biologically independent mice). Data are means with individual data points ±S.E.M. Statistical significance was determined using unpaired two-tailed Student’s *t* test (A, B, and J-M) and two-way ANOVA (C, E, and F-I).

We extended our analysis to young beige APCs to examine the impact of Pdgfrβ activation on IL-33 secretion. Consistent with our hypothesis, Pdgfrβ^D849V^ beige APCs exhibited reduced IL-33 levels in culture media compared to young controls, indicating that Pdgfrβ signaling modulates IL-33 bioavailability (Figure 4C). In agreement, activating Pdgfrβ via Pdgf-BB treatment blunted IL-33 secretion within the media (Figure 4D). We then focused on elucidating the mechanistic link between Pdgfrβ signaling and IL-33 regulation, particularly the role of Stat1[40]. Our previous data showed that Stat1 mediates Pdgfrβ signaling in aged beige APCs[22]. To test this, we inhibited Stat1 using fludarabine in aged SV cells from Pdgfrβ^D849V^ mice in vitro[41]. Remarkably, fludarabine treatment significantly elevated IL-33 secretion from these cells (Figure 4E).

IL-33 regulates the recruitment and activation of ILC2s[42–44]. Therefore, we evaluated if changes in IL-33 expression correlated with ILC2 recruitment. To do so, we FACS isolated CD45+/Lin-/CD25+/CD127+ cells from iWAT and gWAT depots from control and Pdgfrβ-KO aged mice maintained at RT or CE for seven days (Supplementary Fig 4A-C). We confirmed that these were ILC2s by GATA3 and IL-33R staining[45; 46] (Supplementary Fig. 4D, E). In response to Pdgfrβ deletion, we observed more ILC2 cells within iWAT and gWAT depots of aged male and female mice (Fig. 4F-I and Supplementary Fig. 4F, G). We then evaluated if Pdgfrβ activation in young mice prevented ILC2 recruitment. We FACS isolated ILC2s and found that activating Pdgfrβ in young iWAT and gWAT depots prevented ILC2 recruitment in response to cold temperature challenge (Figure 4J-M and Supplementary Fig. 4H, I). Altogether, it appears that Pdgfrβ signaling negatively IL-33 expression to stifle ILC2 accrual.

### Diet-Induced Obesity Impairs Beige Fat Biogenesis and Elevates Pdgfrβ Signaling

Obesity prevents the formation of beige adipocytes[24; 47-49]. We queried whether Pdgfrβ signaling was involved in diet induced obesity-associated beige fat decline. To examine the impact of diet-induced obesity on beige fat biogenesis, we subjected our genetic background control mice (C57BL6/J/129SV) to either a chow or HFD for four and eight weeks, followed by a seven-day CE protocol (Figure 5A). Notably, both male and female mice gain weight on the HFD over the 8-week period (Supplementary Fig. 5A, B). At the four-week mark, both chow and HFD mice exhibited no significant differences in their ability to maintain body temperature during CE (Supplementary Fig. 5C, D). However, after eight weeks on HFD, male and female mice displayed a compromised ability to defend their body temperature during CE (Figure 5B, C). At four weeks of diet, adiposity levels were also comparable between chow and HFD after cold exposure whereas at eight weeks, HFD mice remained elevated (Figure 5D-G). Histological examination using H&E staining and Ucp1 immunostaining revealed a similar presence of Ucp1+ multilocular beige adipocytes within the dorsal lumbar region of iWAT depots across both dietary conditions after four-weeks. However, after eight weeks of HFD, multilocular Ucp1+ beige adipocytes were scarce (Figure 5H-K). This was further supported by Ucp1 quantification revealing a reduction in beige adipocyte biogenesis (Supplementary Fig. 5E, F).

**Figure 5:**
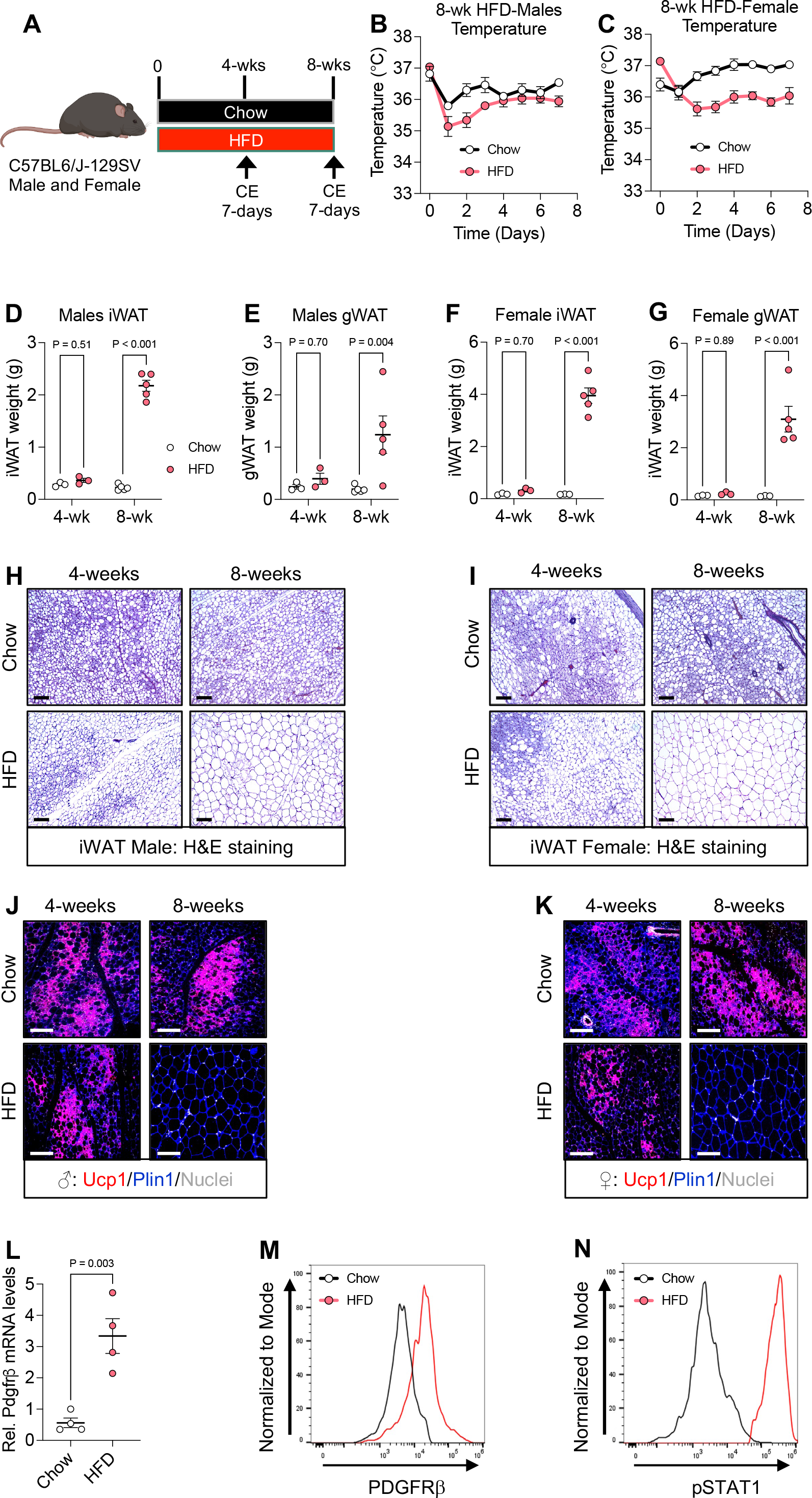
HFD prevents beige fat biogenesis and promotes Pdgfrβ expression. **A.** Experimental design: C57Bl6/J-129SV mice were fed a chow or HFD diet for 4 or 8 weeks. At denoted times mice were subsequently cold exposed for seven days and phenotypically evaluated. **B, C.** Rectal temperature of male (B) and female (C) chow or HFD fed mice throughout CE (n = 3-5 biologically independent mice). **D, E.** iWAT (D) and gWAT (E) weight from chow and HFD male mice (n = 3-5 biologically independent mice). **F, G.** iWAT (F) and gWAT (G) weight from chow and HFD female mice (n = 3-5 biologically independent mice). **H, I.** Representative images of H&E staining of iWAT sections from male (H) or female (I) chow and HFD fed mice at 4 and 8 weeks after CE. **J, K.** Representative images of Ucp1 immunostaining of iWAT sections from male (J) or female (K) chow and HFD fed mice at 4 and 8 weeks after CE. **L.** Relative mRNA levels of Pdgfrβ expression within iWAT depots from male mice fed chow or HFD for 8 weeks maintained at room temperature (RT) (n = 4 biologically independent mice). **M.** Representative flow cytometric histograms of PDGFRβ expression within SMA-positive cells from male mice fed chow or HFD for 8 weeks maintained at RT. **N.** Representative flow cytometric histograms of phosphorylated STAT1 within SMA-positive cells from male mice fed chow or HFD for 8 weeks maintained at RT. Data are means with individual data points ±S.E.M. Statistical significance was determined using unpaired two-tailed Student’s *t* test (B-G and L). Scale bar = 100 µm.

Next, we tested if Pdgfrβ expression was induced by HFD. Indeed, we observed by qPCR analysis that Pdgfrβ was elevated in iWAT depot of HFD male mice (Figure 5L). Moreover, flow cytometric analysis revealed an increase in PDGFRβ protein levels on beige APCs (SMA+) (Figure 5M). Furthermore, we found that STAT1 phosphorylation was also elevated in SMA+ beige APCs after eight-weeks of HFD (Figure 5N). Overall, diet-induced obesity appears to block beige fat formation and lead to the upregulation of Pdgfrβ signaling.

### Restoration of Cold-Induced Beige Adipogenesis in HFD Mice via Pdgfrβ Deletion

To determine whether Pdgfrβ deletion could rescue beige fat biogenesis in the context of HFD, we fed control and Pdgfrβ-KO male and female mice a HFD for eight weeks, followed by TMX administration and a subsequent seven-day CE protocol (Figure 6A). Pdgfrβ-KO male and female mice on HFD maintained homeothermic control more effectively than their HFD control counterparts during CE (Figure 6B, C). However, both groups showed similar cold-induced reductions in body weight (Supplementary Fig. 6A, B). Yet, Pdgfrβ deletion improved glucose tolerance tests, suggesting enhanced glucose clearance in response to CE (Figure 6D, E). In agreement, we observed improved serum triglycerides levels in cold exposed HFD fed Pdgfrβ-KO male and female mice (Supplementary Fig. 6C, D).

**Figure 6:**
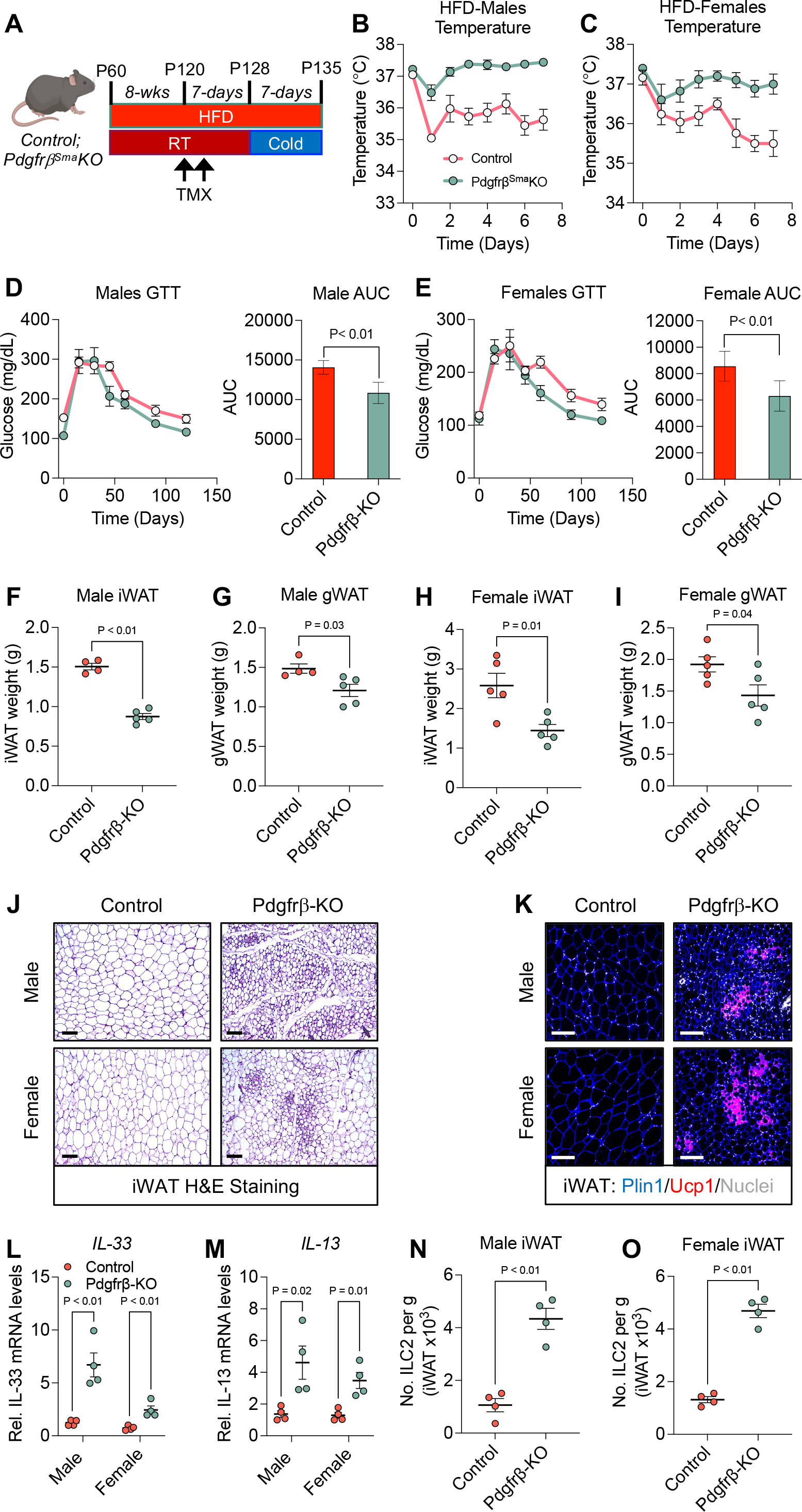
Deleting Pdgfrβ restores cold temperature induced metabolic parameters and beige fat development in diet induced mice. **A.** Experimental design to evaluate Control and Pdgfrβ-KO diet induced obese male and female mice. Mice were fed a HFD for 8 weeks and subsequently administered TMX to induce Pdgfrβ deletion. After a seven-day TMX washout, mice were exposed to cold temperatures for seven days. **B, C.** Rectal temperature of male (B) and female (C) Control and Pdgfrβ-KO aged mice throughout CE (n = 4-5 biologically independent mice). **D.** Intraperitoneal glucose tolerance test performed on Control and Pdgfrβ-KO obese male mice after seven days of CE. Right: Area under the curve (AUC) was calculated (n = 4-5 biologically independent mice). **E.** Intraperitoneal glucose tolerance test performed on Control and Pdgfrβ-KO aged female mice after seven days of CE (n = 5 biologically independent mice). Right: Area under the curve (AUC) was calculated. **F, G.** iWAT (F) and gWAT (G) weights from Control and Pdgfrβ-KO obese male mice after seven days of CE (n = 4-5 biologically independent mice). **H, I.** iWAT (H) and gWAT (I) weights from Control and Pdgfrβ-KO obese female mice after seven days of CE (n = 5 biologically independent mice). **J.** Representative images of H&E staining of iWAT sections from Control and Pdgfrβ-KO male and female obese mice maintained at CE for seven days. **K.** Representative images of Ucp1 immunostaining of iWAT sections from Control and Pdgfrβ-KO male and female obese mice maintained at CE for seven days. **L.** Relative mRNA levels of IL-33 within iWAT depots from cold exposed Control and Pdgfrβ-KO male and female obese mice (n = 4 biologically independent mice). **M.** Relative mRNA levels of IL-13 within iWAT depots from cold exposed Control and Pdgfrβ-KO male and female obese mice (n = 4 biologically independent mice). **N, O.** Relative abundance of ILC2s within iWAT depots of male (N) and female (O) cold exposed Control and Pdgfrβ-KO male and female obese mice (n = 4 biologically independent mice). Data are means with individual data points ±S.E.M. Statistical significance was determined using unpaired two-tailed Student’s *t* test (B, C, and F-K). Scale bar = 100 µm.

In line with beige adipogenesis, Pdgfrβ-KO mice exhibited a significant reduction in adiposity compared to HFD controls (Figure 6F-I). However, non-adipose tissue weight remained comparable between control and mutant mice (Supplementary Fig. 6E, F). Histological analysis of iWAT sections from Pdgfrβ-KO mice revealed the presence of smaller white adipocytes and the emergence of multilocular beige adipocytes (Figure 6J and Supplementary Fig. 6G, H). Immunostaining for Ucp1 confirmed robust beige adipocyte formation in the iWAT of Pdgfrβ-KO mice (Figure 6K and Supplementary Fig. 6I, J). Although gWAT depots were reduced in size, beige fat development was not detected post-CE (Supplementary Fig. 6K, L). Supporting these histological findings, qPCR analyses demonstrated a restoration of thermogenic gene expression in iWAT samples from HFD-fed Pdgfrβ-KO mice (Supplementary Fig. 6M, N). Together, these results indicate that Pdgfrβ deletion effectively reinstates the ability to generate beige adipocytes in response to cold exposure, even under conditions of diet-induced obesity.

IL-33 bioavailability and ILC2 recruitment and activation are reduced in obese mice and humans[50–54]. Therefore, we investigated if Pdgfrβ deletion reinstated IL-33 expression and ILC2 accrual and activation. Indeed, we observed that Pdgfrβ deletion elevated IL-33 expression within iWAT depots of male and female mice after CE (Figure 6L). In addition, we observed higher levels of IL-13, a byproduct of ILC2 activation[44; 55] (Figure 6M). Furthermore, FACS analysis revealed more ILC2s within iWAT depots of Pdgfrβ-KO male and female mice (Figure 6N, O). Consistent with Pdgfrβ deletion, we observed less STAT1 phosphorylation in male and female iWAT depots, suggesting that Stat1 may mediate Pdgfrβ signaling in HFD administered mice (Supplementary Fig. 6O). Taken together, it appears that HFD invokes Pdgfrβ upregulation and signaling to prevent IL-33 expression and ILC2 accrual and activation thereby blunting beige fat development.

## Discussion

Cold temperatures stimulate the formation of beige adipocytes to coordinate thermogenesis and metabolism. The appearance of beige adipocytes upsurges metabolic demand by increasing energy expenditure, shrinking adipose tissues, and lowering serum lipid levels[5; 28; 29; 56-58]. This effect is considered protective against obesogenic cues. However, the ability to generate beige fat declines with age and obese conditions, a time when beige fat would be the most valuable[4; 20]. While aging and obesity are associated with an increased risk of metabolic dysfunction, the underlying mechanisms of beige fat decline remain complex. This study highlights the pivotal role of Pdgfrβ signaling in the regulation of beige APC function and metabolic health, particularly in the contexts of aging and obesity. Our findings demonstrate that deletion of Pdgfrβ in aged mice not only restores the formation of metabolically functional beige adipocytes but also enhances energy expenditure, improves glucose tolerance, and promotes lipid oxidation. These metabolic improvements underscore the potential of targeting Pdgfrβ signaling to counteract the detrimental effects of aging and obesity on metabolic health.

Interestingly, cold exposure (CE) has been shown to exert metabolic benefits beyond beige fat development[30]. Our data reveal that CE effectively reduces body weight, enhances glucose metabolism, improves blood lipid profiles, and decreases adiposity in both young and aged mice, independent of beige fat formation. This suggests that the metabolic demand induced by CE alone is sufficient to drive favorable metabolic changes. However, the precise tissues and mechanisms mediating these effects remain to be elucidated. It is likely that multiple tissues orchestrate responses to meet the energy demands induced by CE, thereby maintaining homeostasis under colder environmental conditions. Human tracer studies have provided insights into the tissues and substrates responsive to cold exposure[59–61]. Notably, in humans, skeletal muscle may play a more significant role in thermogenic and metabolic responses than adipose tissue[60]. Continued research into substrate utilization and tissue uptake during CE will enhance our understanding of how different tissues contribute to the metabolic adaptations observed under cold exposure, including the role of thermogenic fat cells.

The interplay between Pdgfrβ signaling and the immune niche is particularly intriguing to beige adipocyte regulation. Our study associates Pdgfrβ activation with reduced IL-33 secretion from beige adipocyte progenitors, leading to decreased ILC2 accrual and activation. This immunological shift appears to be a key mechanism by which Pdgfrβ signaling impedes beige fat formation. Despite the critical role of these signals in coordinating beige fat development, the specific cellular sources or pools of beige adipocytes remain unidentified. Our previous fate mapping analysis revealed that beige adipocyte progenitor cells (APCs) remain senescent following Pdgfrβ deletion, raising questions about the cell types involved in generating beige adipocytes without Pdgfrβ signaling. Furthermore, the role of ILC2 cells in inducing beige fat development warrants further investigation. Although methionine-enkephalin, an ILC2 bioproduct, has been implicated in promoting beige fat formation, the underlying mechanisms remain undefined[50; 62]. Broadly, dietary and age-associated changes in WAT are accompanied by immune cell composition modifications[63]. Whether Pdgfrβ regulates a more diverse immunological cell composition remains to be explored.

Our findings also shed light on the impact of diet-induced obesity on beige fat biogenesis, paralleling the effects of aging. Mice fed a high-fat diet (HFD) exhibited reduced thermogenic capacity and increased adiposity, alongside upregulated Pdgfrβ signaling. Remarkably, Pdgfrβ deletion reinstated beige adipocyte generation in response to CE, even in the obese state. These interventions led to improved thermogenesis, enhanced glucose metabolism, and reduced adiposity, highlighting the therapeutic potential of targeting Pdgfrβ signaling in metabolic diseases. However, a deeper understanding of the transcriptional regulation of Pdgfrβ in beige APC biology is essential. Elucidating the upstream mechanisms governing Pdgfrβ will uncover the molecular underpinnings driving beige APC dysfunction and niche regulatory signals. Interestingly, several studies have indicated that perivascular cells express and produce IL-33[64; 65]. Moreover, beta-adrenergic receptor activation promotes gene transcription of IL-33 within Pdgfrβ+ APCs[62; 66]. However, the molecular mediators regulating the age-dependent upregulation of Pdgfrβ expression to limit IL-33 transcription remains to be define.

In summary, our study emphasizes the critical role of Pdgfrβ in regulating beige fat progenitor function and metabolic health. By modulating Pdgfrβ signaling, we can restore the beneficial effects of beige fat biogenesis and mitigate the metabolic detriments associated with aging and obesity. These findings open new avenues for therapeutic strategies aimed at enhancing beige fat formation and improving metabolic outcomes in vulnerable populations. Future research should focus on identifying the precise cellular and molecular mechanisms underlying these processes to fully harness the potential of targeting Pdgfrβ signaling in metabolic health interventions.

## Acknowledgements

The authors thank the Berry lab help advice and discussions, specifically Caila Low and Derek Lee. The authors thank the Cornell Biotechnology Resources Center Flow Cytometric Core Facility and the Center of Animal Resources and Education for excellent assistance with experimental collection and mouse husbandry, respectively.

## Data availability

The data that support the findings of this study are available from the corresponding author (DCB) upon reasonable request.

## Author Contributions

A.M.B. and D.C.B conceived and designed the study and wrote the manuscript. A.M.B performed all the experiments, collected data, and analyzed the results.

## METHODS

### MOUSE MODELS

All animal experiments were performed according to the procedures approved by the Cornell University Institutional Animal Care and Use Committee under the auspices of protocol number 2017-0063. The Sma-Cre^ERT2^ mouse model was obtained from Drs. Pierre Chambon and Daniel Metzger[32]. Sma-Cre^ERT2^ mice were then crossed with either Pdgfrβ^D849V^ (stock #018435[37]) or Pdgfrβ^fl/fl^ (stock #010977[67]) purchased from the Jackson Laboratories. Prior to experimentation, offspring were intercrossed for six generations and maintained on a mixed C57BL6/J-129SV background. Mice were maintained on a 14:10-hour light/dark cycle with free access to food and water. To induce recombination, the denoted mice were administered one dose of TMX (50 mg/Kg; Cayman Chemical: 13258) dissolved in sunflower seed oil (Sigma, item no: S5007) for two consecutive days via intraperitoneal (IP) injection. Post the final TMX injection, mice were maintained for seven days at room temperature (RT, ∼25°C) as a TMX washout period. For cold temperature exposure, mice seven days post TMX were housed in a 6.5°C cold chamber (Power Scientific RIS70SD). All animal experiments were performed on 3 or more mice per cohort and performed at least twice. All animal experiments were performed according to procedures approved by the Cornell University Institutional Animal Care and Use Committee under the auspices of protocol number 2017-0063.

### PHYSIOLOGICAL MEASUREMENTS

Rectal temperatures were monitored daily (∼18:00 EST) using a TH-5 Thermalert Clinical Thermometer (Physitemp) attached to a RET-3 rectal probe for mice (Physitemp). Prior to insertion, the probe was lubricated with glycerol then inserted 1.27 centimeters (1/2 inch). Temperatures were recorded once the reading stabilized.

### GLUCOSE TOLERANCE TEST

Glucose tolerance tests were performed on mice fasted for 6 hours at room temperature. Random glucose was measured prior to fasting. Tail blood glucose levels were first measured at 0 (fasted) then mice were then i.p. injected with glucose (1.25g/kg; Sigma G8270, dissolved in sterile water). Tail blood glucose levels were then measured at 15, 30, 45, 60, 90- and 120-mins post injection with a Bayer Contour glucometer and strips.

### METABOLIC CAGE ANALYSIS

All mice were singly housed and acclimated for two days at RT in a Promethion metabolic screening system prior to data collection. All cages were equipped with food hoppers, water bottles, and shelter huts for continuous food and water intake and body weight monitoring. Cages were housed in an environmentally controlled cabinet, measuring oxygen consumption, carbon dioxide production, and total locomotor activity to calculate the respiratory exchange ratio and total energy expenditure. After the acclimation period, mice were then out of the cages, the recording paused, and the chamber chilled to 6.5 °C for cold temperature measurements. Mice were then kept at 6.5 °C and the data recorded for 7 days in the cold. Data analysis was conducted using the web based indirect calorimetry software CalR (Mina et al., 2018).

### HIGH-FAT DIET

Mice were fed a high fat diet (Research diets; D12492; 60% fat) for either 4 or 8 weeks prior to TMX injection and cold exposure. Prior to and at the completion of the diet body weights were collected.

### BLOOD CHEMISTRY

Blood samples were obtained via heart puncture and left to clot for 2 hours at room temperature. Subsequently, the samples were centrifuged at 7500 rpm for 5 minutes to separate serum, which was then aliquoted and frozen. Serum triglyceride levels were determined using the Triglyceride Quantification Colorimetric/Fluorometric Kit (Sigma-Aldrich Catalog: MAK266) from Millipore-Sigma as per the manufacturer’s instructions, with a 1:25 serum dilution before assessment. Serum cholesterol levels were measured with the Cholesterol Quantification Kit (Sigma-Aldrich Catalog: MAK043) from Millipore-Sigma, following a 1:25 serum dilution. Lastly, Adiponectin serum concentrations were quantified using the Quantikine ELISA Mouse Adiponectin/Acrp30 (Catalog: MRP30) from R&D Systems in accordance with the manufacturer’s guidelines after a 1:2000 serum dilution.

### ADIPOSE SV CELL ISOLATION

Adipose tissues were removed from a single mouse, and the inguinal or visceral depots were finely diced before being immersed in 10 ml of isolation buffer composed of 0.1 M HEPES, 0.12 M NaCl, 50 mM KCl, 5 mM D-glucose, 1.5% BSA, and 1 mM CaCl_2_, supplemented with collagenase type I (10,000 units). Samples were then subjected to incubation in a 37°C incubator with gentle agitation for approximately 1 hour. Following digestion, the tissue was combined with serum-free Dulbecco’s Modified Eagle’s Medium Nutrient Mixture F-12 Ham (Sigma, cat. no. D8900 and D6421) (DMEM/F12) and filtered through a 100 (iWAT) or 70 (gWAT) µm cell strainer. Samples were then centrifuged at 1250 rpm for 10 minutes, after which the supernatant was discarded, and the remaining pellet resuspended in 10 ml of erythrocyte lysis buffer containing 155 mM NH_4_Cl, 10 mM KHCO_3_, and 0.1 mM EDTA. Following a 5-minute incubation period, growth media (DMEM/F12 with 10% fetal bovine serum (FBS)) was added, mixed and stained through a 40 µm cell strainer. The sample was then centrifuged at 1250 rpm for 5 minutes, the supernatant removed, and the pellet resuspended in growth media for subsequent plating. Alternatively, cells were harvested for flow cytometric analysis.

### IL-33 ELISA

The iWAT SVF was isolated as described above, and cells were plated in a 96-well plate in growth media. Growth media was changed the following morning (∼18 hours) post plating. IL-33 was measured according to the manufactures protocol for cell culture samples. Aliquots of cell culture supernates were collected and spun at 500 x g for 5 minutes to remove particulates, per the manufacture’s suggestion. 50 µl of sample was then assayed immediately in duplicate and the absorbance measured.

### FLOW CYTOMETRY

The iWAT or gWAT SVF was isolated as described above and resuspended in 1X PBS along with blue fluorescent reactive dye. Cells were then pelleted and resuspended in 0.3 −0.5 ml of FACS buffer consisting of 2.5% horse serum; 2 mM EDTA in 1X PBS with 1X protease/phosphatase inhibitor cocktail in and filtered through a 5 ml cell-strainer capped FACS tube (BD Falcon). Cell sorting was performed on BD Biosciences FACS Aria Fusion. Alternatively, cells were analyzed on a Thermo-Fisher Attune NxT cytometry. Viable cells were gated from the blue fluorescent reactive dye negative population followed by singlet forward and side scatter pattern. Cells were fixed for 30 minutes in 4% paraformaldehyde on ice, permeabilized, stained and analyzed for phosphorylated Stat1 (1:200; 9167S Cell Signaling) or total Pdgfrβ (1:200; 4564S Cell Signaling) conjugated to Cy5 donkey anti-rabbit secondary antibody (1:200; A10523 Invitrogen). Alternatively, CD45+ (1:200; 103151 Biolegend) cells were FACS isolated from the total iWAT or gWAT SV cell population of respective mice. For ILC2 analysis, live CD45+ cells were stained with the anti-mouse lineage negative cocktail (1:200; 133303 Biolegend), CD25 (1:250; 135019 Biolegend), and CD127 (1:250; 101907 Biolegend) antibodies and analyzed for their respective conjugated fluorophore. For ILC2 validation, cells were additionally stained for either Gata3 (1:200; 12-9966-41 Invitrogen) or IL33R (1:200; 566311 BD Biosciences) and analyzed for their respective conjugated fluorophore.

### RNA ISOLATION AND qPCR

Approximately 200 mg of WAT or the quadricep muscle from a single mouse was placed into Precellys tubes containing ceramic (WAT) or metal (skeletal muscle) beads and 1 ml of TRIzol reagent (ABI). Tissues were then homogenized in a Precellys 24 homogenizer using the following the settings: 3 pulses at 4500 rpm for 30 seconds with a 30 second rest between pulses with a final rest of 4 minutes. RNA was isolated through conventional chloroform extraction and isopropanol precipitation technique. The concentration and quality of RNA were measured using a TECAN infinite F-nano+ spectrophotometer. Subsequently, 1 µg of RNA was reverse transcribed into complementary DNA (cDNA) utilizing the high-capacity RNA to cDNA kit (Life Technologies #4368813). For qPCR, cDNA was diluted 1:10 and added to PowerUp™ SYBR™ Green Master Mix (Life Technologies A25742) along with denoted primers (Table S1). qPCR analysis was performed on an Applied Biosystems QuantStudio™ 3 Real-Time PCR System using the DD-CT method compared to the internal control, Rn18s. Data points correspond to individual biological replicates, with each data point assessed in quadruplicate technically. Details regarding the qPCR primer sequences can be accessed in Table S1.

### HISTOLOGICAL ANALYSIS

Adipose tissues were dissected, weighed and immediately placed in 10% formalin (neutralized with 1X PBS) for 24 hours. Tissues were then processed using Thermo Scientific™ STP 120 Spin Tissue Processor with the following conditions: Bucket 1: 50% ethanol (45 minutes); Bucket 2: 70% ethanol (45 minutes) Bucket 3: 80% ethanol (45 minutes); Bucket 4 and 5: 95% ethanol (45 minutes); Bucket 6 and 7: 100% ethanol (45 minutes); Bucket 8-10: Xylene Substitute (45 minutes); Bucket 11 and 12: paraffin (4 hours each). A Histostar™ embedding station was used to embed tissues into cassettes using and blocks were refrigerated for at least 24 hours prior to sectioning. 8-12-micron tissue sections were generated using a HM-325 microtome using low profile blades. Sections were placed in a 40 °C water bath and positioned on microscope slides. Slides were then baked overnight at 55 °C prior to staining.

Hematoxylin and eosin (H&E) staining: Slides were rehydrated using the following protocol: xylene (3 mins 3X), 100% reagent alcohol (1 min 2X); 95% reagent alcohol (1 min 2X); water (1 min). Slides were stained in hematoxylin for 2 minutes 30 seconds followed by 10 repeated submerges in eosin staining. Slides were then dehydrated in the reverse order of the above rehydration steps. Slides were then cover slipped with Cryoseal 60 mounting media. Brightfield images were acquired using a Leica DMi8 inverted microscope system.

Immunofluorescent staining: Slides were rehydrated as described above. Antigen retrieval was performed using 1X Citrate Buffer, made from 10X stock (Electron Microscopy Sciences R-Buffer A, 10X, pH 6; Catalog: 62706-10) diluted in diH_2_O, and placed into an antigen retriever pressure cooker (EMS Catalog #62706) for 2 hours. Slides were then permeabilized using 0.3%TritonX-100 in 1X TBS for 30 minutes and washed three times in 1X TBS. Slides were then blocked with 10% donkey serum in 1X TBS for 30-minutes and incubated with primary antibody diluted in blocking solution overnight at 4°C. The following primary antibodies were used: guinea pig anti-perilipin (1:200; Fitzgerald 20R-PP004); rabbit anti-Ucp1 (1/200; abcam: ab10983); rat anti-F4/80 (1:200; R&D Systems MAB5580-SP). Secondary antibodies from Invitrogen or Jackson ImmunoResearch were used at 1/200 dilutions and incubated for 2 hours at room temperature. The following secondary antibodies were used: Cy5 donkey anti-guinea pig, Texas red donkey anti-rabbit, or 488 donkey anti-rat. Slides were washed and stained with Hoechst (H3570; Life Technologies) (1 µg/ml in 1X TBS) for 10 minutes and cover slipped with Thermo Scientific™ Shandon™ Immu-Mount™ mounting media. Fluorescent images were collected on a Leica DMi8 inverted microscope system.

### ADIPOCYTE AREA QUANTIFICATION

Three 10X images were assessed from at least three mice/cohort for adipocyte area quantification. The Fiji ImageJ Adiposoft plugin was used to calculate adipocyte area; adipocyte areas below 50 microns were omitted due to reliability. NIH Fiji Image J software was additionally used to quantify fibrotic tissue area.

### BEIGE AREA QUANTIFICATION

Three 20X immunofluorescent images containing the individual channel for either Ucp1 or F4/80 from 3 mice per group were quantified for the percent of beige or F4/80 positive area utilizing ImageJ Fiji software. Images were converted to the appropriate color threshold and highlighted to determine percent fluorescence over total fluorescent signal. Output values were then corrected for background signal and averaged in their respective groups.

### QUANTIFICATION AND STATISTICAL ANALYSIS

Statistical significance was assessed by either one- or two-way ANOVA for two-group or multiple group comparisons. Data are means and error bars are expressed as ± SEM. A P value of < 0.05 was considered significant in all experiments. The statistical parameters and the number of mice used per experiment are found in the figure legends. Mouse experiments were performed in biological duplicate or triplicate with at least three mice per cohort. The Leica Application suite X Microscope software was used for image acquisition and analyses. The NIH Fiji Image J software was used to quantify adipocyte, fibrotic, Ucp1+, and F4/80+ areas. For all experiments, three random fields were assessed from at least three mice/cohort. The flow cytometric analysis software, FlowJo, was used to analyze cell populations, antibody staining, and generate flow cytometric plots. All graphs and statistical analyses were performed using GraphPad Prism 9. Excel was used for raw data collection, analysis, and quantification.

