## Supplementary Figures for "Reversing Pdgfrβ Signaling Restores Metabolically Active Beige Adipocytes by Alleviating ILC2 Suppression in Aged and Obese Mice"

Supplementary Figure 1

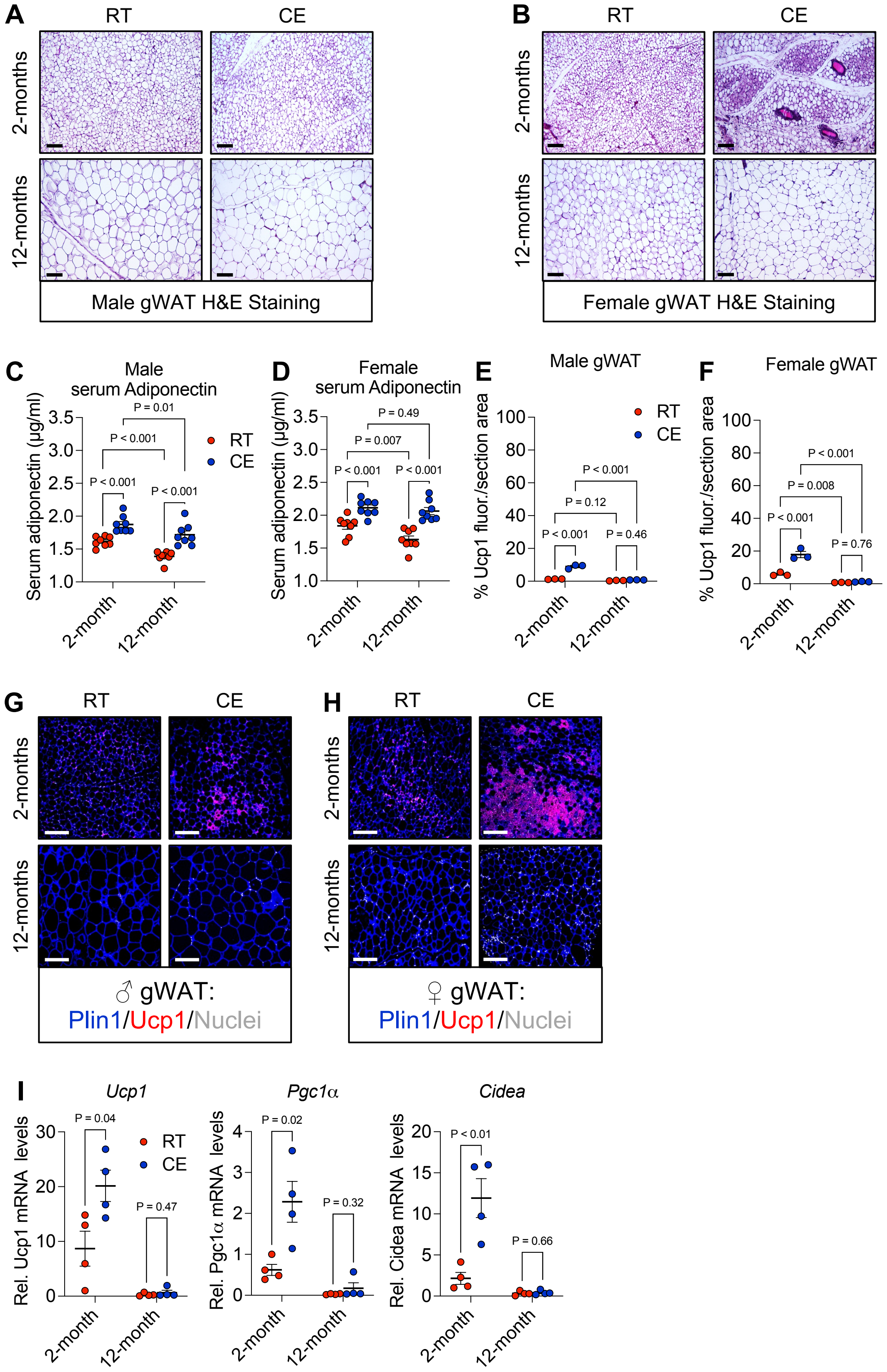

**Supplementary Figure 1: Cold exposure improves metabolic parameters in aged mice, independent of beige fat formation.**

**A.** Representative images of H&E staining of gWAT sections from male mice maintained at RT or CE at 2 or 12 months of age.

**B.** Representative images of H&E staining of gWAT sections from male mice maintained at RT or CE at 2 or 12 months of age.

**C, D.** Serum adiponectin levels in male (C) and female (D) mice maintained at RT or CE at 2 or 12 months (n = 8 biologically independent mice).

**E, F.** Quantification of Ucp1 immunostaining of gWAT sections from male (E) and female (F) mice maintained at RT or CE at 2 or 12 months of age (n = 3 biologically independent mice).

**G, H.** Representative images of H&E staining of gWAT sections from male (G) and female (H) mice maintained at RT or CE at 2 or 12 months of age.

**I.** Relative mRNA levels of denoted thermogenic genes within iWAT depots from male mice maintained at RT or CE at 2 or 12 months of age (n = 4 biologically independent mice).

Data are means with individual data points ±S.E.M. Statistical significance was determined using one-way ANOVA (C-F) and unpaired two-tailed Student's *t* test (I). Scale bar = 100 μm.

Supplementary Figure 2

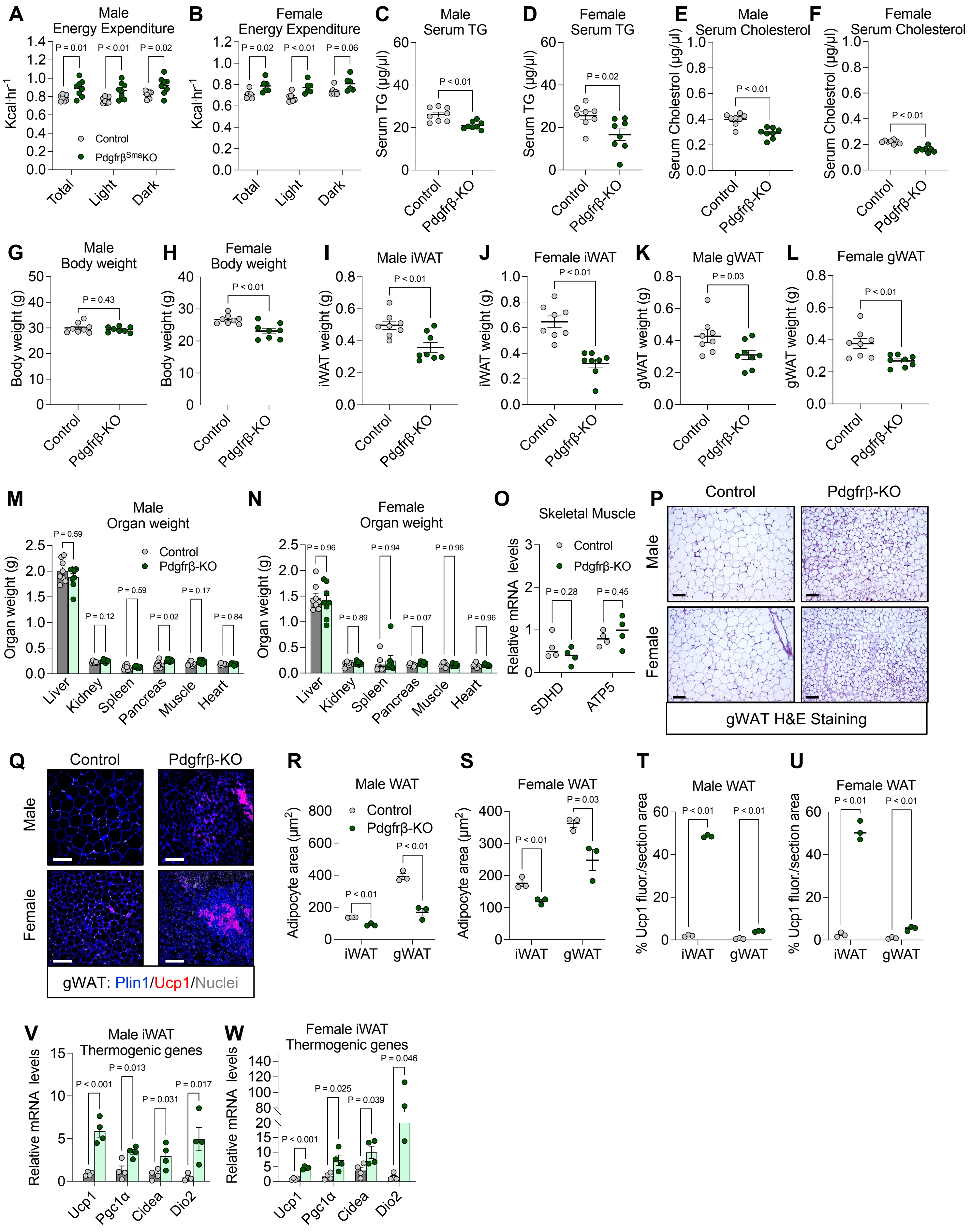

**Supplementary Figure 2: Deletion of Pdgfrβ improves cold temperature induced metabolic parameters and beige fat in aged mice.**

**A, B.** Averaged energy expenditure graphs from Control and Pdgfrβ-KO aged male (A) and female (B) mice (n = 6-8 biologically independent mice).

**C, D.** Serum triglyceride (TG) levels from Control and Pdgfrβ-KO aged male (C) and female (D) mice after CE (n = 8 biologically independent mice).

**E, F.** Serum cholesterol levels from Control and Pdgfrβ-KO aged male (E) and female (F) mice after CE (n = 8 biologically independent mice).

**G, H.** Body weight of Control and Pdgfrβ-KO aged male (G) and female (H) mice after CE (n = 8 biologically independent mice).

**I, J.** iWAT weight from Control and Pdgfrβ-KO aged male (I) and female (J) mice after CE (n = 8 biologically independent mice).

**K, L.** gWAT weight from Control and Pdgfrβ-KO aged male (K) and female (L) mice after CE (n = 8 biologically independent mice).

**M, N.** Tissue weight from Control and Pdgfrβ-KO aged male (M) and female (N) mice after CE (n = 8 biologically independent mice).

**O.** Relative mRNA expression levels of skeletal muscle shivering genes (n = 4 biologically independent mice).

**P.** Representative images of H&E staining of gWAT sections from Control and Pdgfrβ-KO male and female mice after CE.

**Q.** Representative images of Ucp1 immunostaining of gWAT sections from Control and Pdgfrβ-KO male and female mice after CE.

**R, S.** Quantification of adipocyte area from iWAT and gWAT section from Control and Pdgfrβ-KO male (R) and female (S) mice after CE (n = 3 biologically independent mice).

**T, U.** Quantification of Ucp1 immunostaining from iWAT and gWAT section from Control and Pdgfrβ-KO male (T) and female (U) mice after CE (n = 3 biologically independent mice).

**V, W.** Relative mRNA expression levels of denoted thermogenic genes within iWAT depots from Control and Pdgfrβ-KO male (V) and female (W) mice after CE (n = 4 biologically independent mice).

Data are means with individual data points ±S.E.M. Statistical significance was determined using unpaired two-tailed Student's *t* test (B-O, and R-W). Scale bar = 100 μm.

Supplementary Figure 3

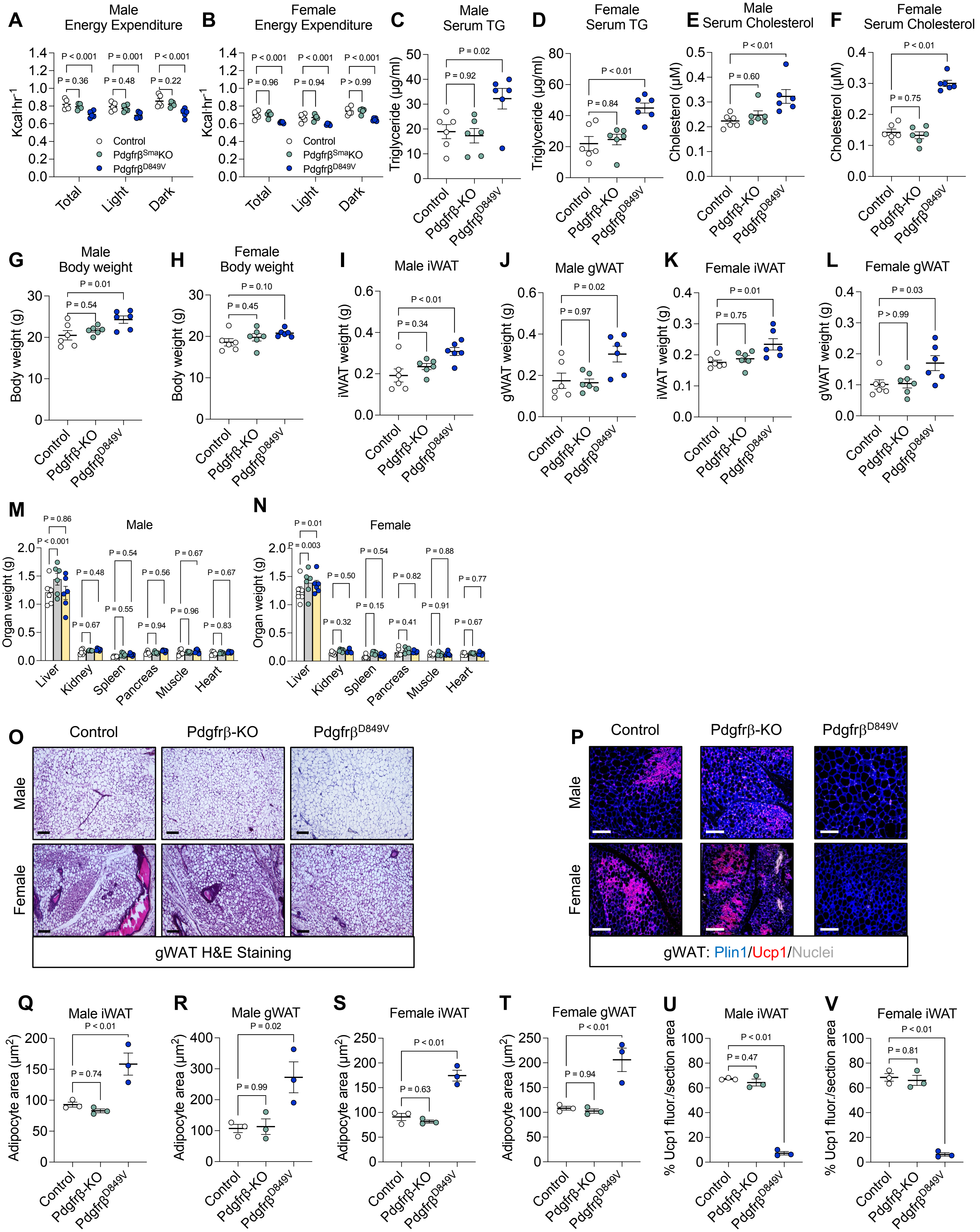

**Supplementary Figure 3: Constitutive activation of Pdgfrβ diminishes cold temperature induced metabolic parameters and blocks beige fat development in young mice.**

**A, B.** Averaged energy expenditure graphs from Control, Pdgfrβ-KO, and Pdgfrβ$^{D849V}$ young (2-month-old) male (A) and female (B) mice after CE (n = 5 biologically independent mice).

**C, D.** Serum triglyceride (TG) levels from Control, Pdgfrβ-KO, and Pdgfrβ$^{D849V}$ young male (C) and female (D) mice after CE (n = 6 biologically independent mice).

**E, F.** Serum cholesterol levels from Control, Pdgfrβ-KO, and Pdgfrβ$^{D849V}$ young male (E) and female (F) mice after CE (n = 6 biologically independent mice).

**G, H.** Body weight of Control, Pdgfrβ-KO, and Pdgfrβ$^{D849V}$ young male (G) and female (H) mice after CE (n = 6 biologically independent mice).

**I, J.** iWAT and gWAT weight from Control, Pdgfrβ-KO, and Pdgfrβ$^{D849V}$ young male mice after CE (n = 6 biologically independent mice).

**K, L.** iWAT and gWAT weight from Control, Pdgfrβ-KO, and Pdgfrβ$^{D849V}$ young female mice after CE (n = 6 biologically independent mice).

**M, N.** Tissue weight from Control, Pdgfrβ-KO, and Pdgfrβ$^{D849V}$ young male (M) and female (N) mice after CE (n = 6 biologically independent mice).

**O.** Representative images of H&E staining of gWAT sections from Control, Pdgfrβ-KO, and Pdgfrβ$^{D849V}$ young male and female mice after CE.

**P.** Representative images of Ucp1 immunostaining of gWAT sections from Control, Pdgfrβ-KO, and Pdgfrβ$^{D849V}$ young male and female mice after CE.

**Q-T.** Quantification of adipocyte area from iWAT and gWAT section from Control, Pdgfrβ-KO, and Pdgfrβ$^{D849V}$ young male (Q,R) and female (S, T) mice after CE (n = 3 biologically independent mice).

**U, V.** Quantification of Ucp1 immunostaining from iWAT section from Control, Pdgfrβ-KO, and Pdgfrβ$^{D849V}$ male (U) and female (V) mice after CE (n = 3 biologically independent mice).

Data are means with individual data points ±S.E.M. Statistical significance was determined using unpaired two-tailed Student's *t* test (B, C, and F-K). Scale bar = 100 μm.

Supplementary Figure 4

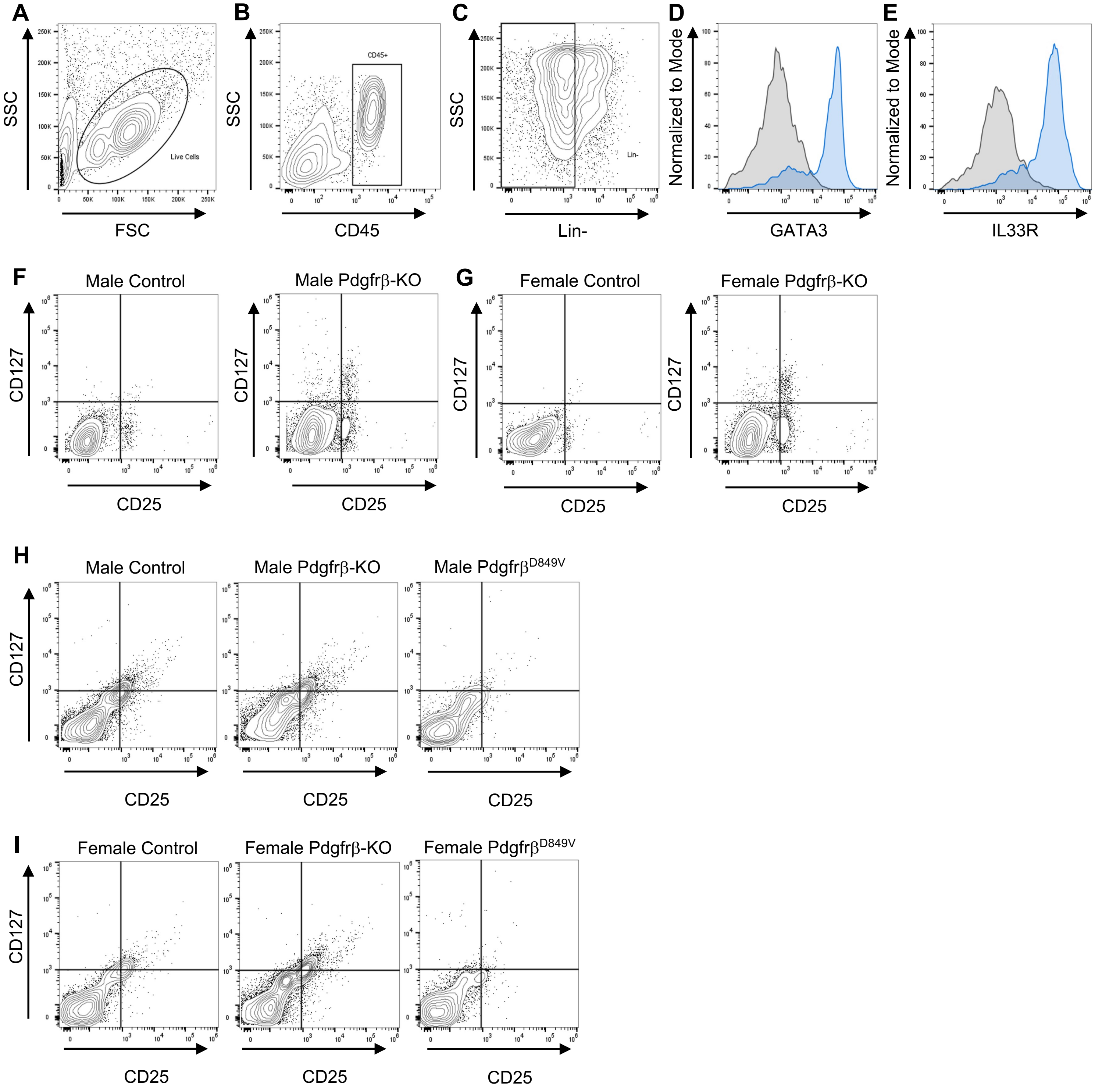

**Supplementary Figure 4: Pdgfrβ associates with IL-33 bioavailability and prevents ILC2 accrual.**
**A.** Representative FACS gating strategy for live cells.
**B.** Representative FACS gating strategy for CD45 cells.
**C.** Representative FACS gating strategy for CD45+/Lin- cells.
**D, E.** Representative FACS histograms of GATA3 and IL-33R for ILC2 confirmation.
**F, G.** Representative FACS plots of ILC2 abundance within iWAT depots from aged Control and Pdgfrβ-KO male (F) and female (G) mice after CE.
**H, I.** Representative FACS plots of ILC2 abundance within iWAT depots from young Control, Pdgfrβ-KO, and Pdgfrβ$^{D849V}$ male (H) and female (I) mice after CE.

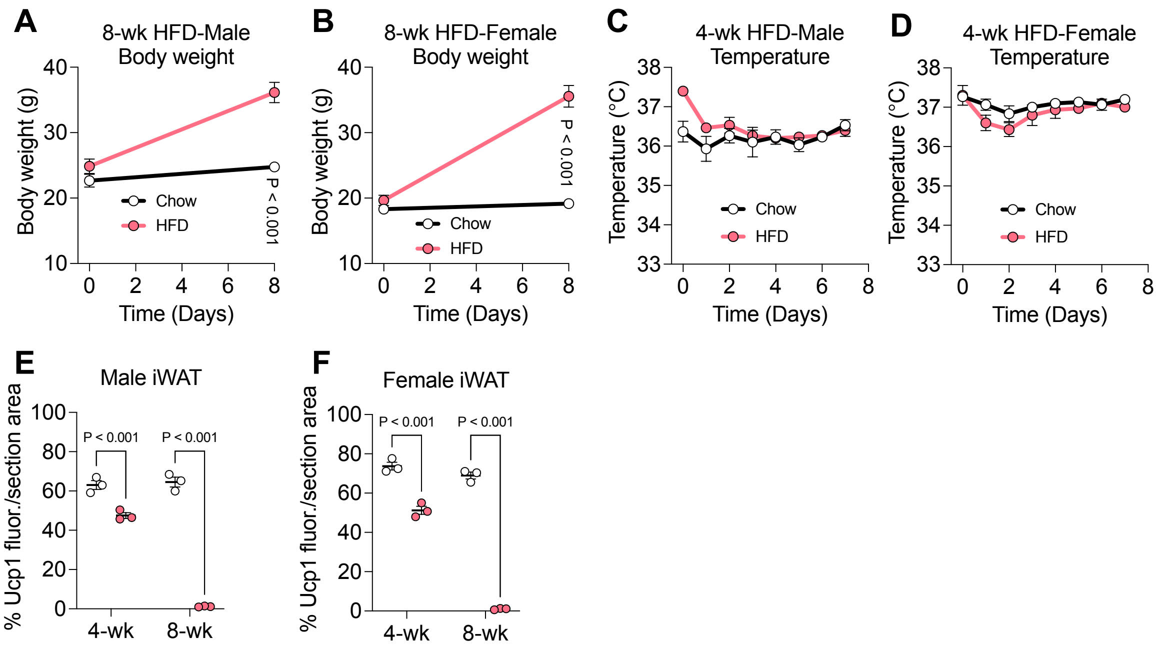


**Supplementary Figure 5: HFD prevents beige fat biogenesis and promotes Pdgfrβ expression.**
**A, B.** Body weight curve of C57Bl6/J-129SV male (A) and female (B) mice fed a chow or HFD diet for eight weeks (n = 3-5 biologically independent mice).
**C, D.** Rectal temperature of male (B) and female (C) chow or HFD fed mice throughout CE after four-weeks of diet (n = 3-5 biologically independent mice).
**E, F.** Quantification of Ucp1 immunostaining from iWAT section from chow and HFD fed male (E) and female (F) mice at four- and eight-weeks after CE (n = 3 biologically independent mice).
Data are means with individual data points ±S.E.M. Statistical significance was determined using unpaired two-tailed Student's *t* test (A-F).

Supplemental Figure 6

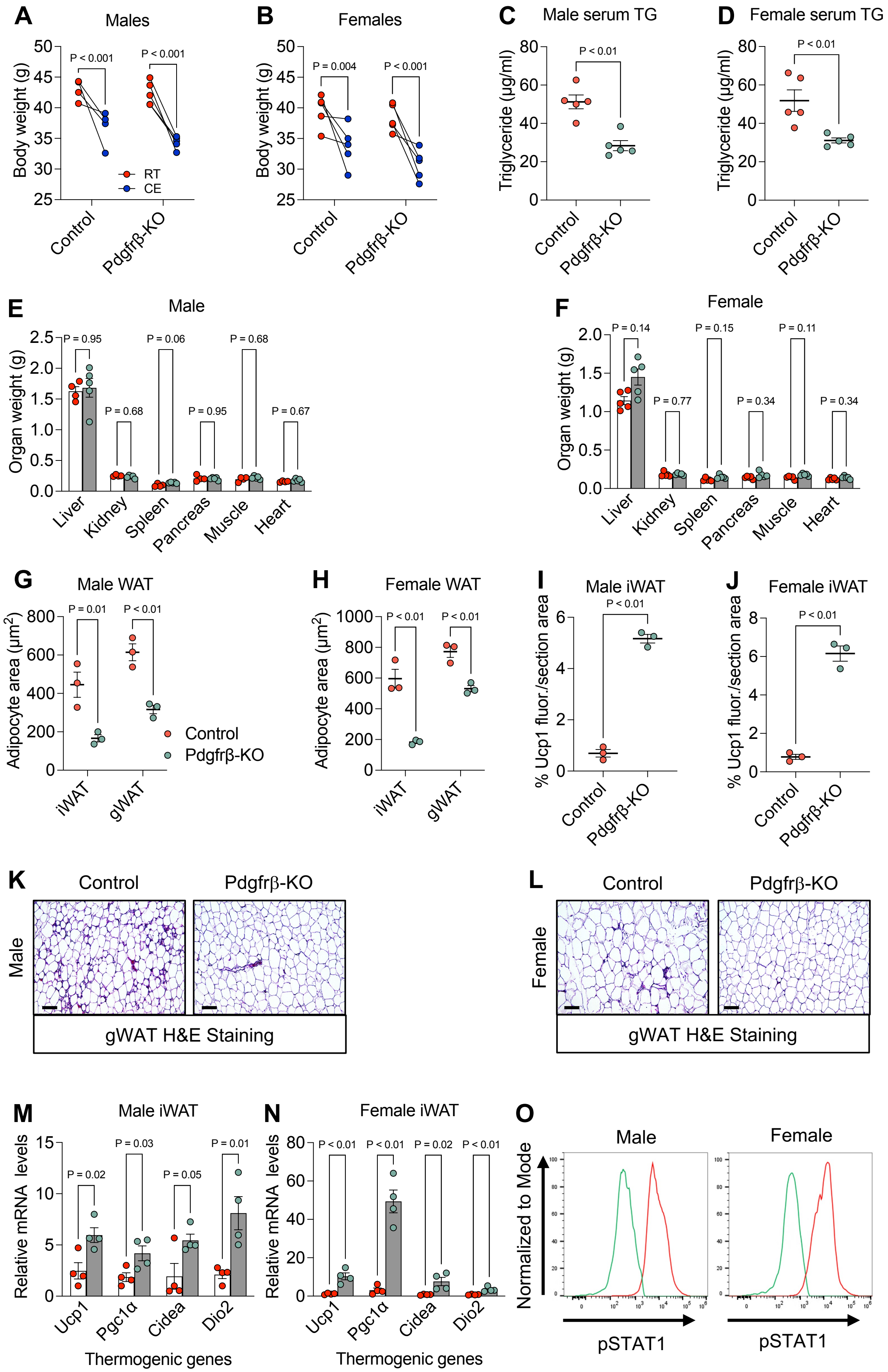

**Supplemental Figure 6: Deleting Pdgfrβ restores cold temperature induced metabolic parameters and beige fat development in diet induced mice.**

**A, B.** Body weights of Control and Pdgfrβ-KO diet induced obese male (A) and female (B) mice before and after seven days of cold temperature exposure (n = 4-5 biologically independent mice).

**C, D.** Serum triglycerides (TG) levels in Control and Pdgfrβ-KO diet induced obese male (C) and female (D) mice after seven days of cold temperature exposure (n = 4-5 biologically independent mice).

**E, F.** Tissue weights from Control and Pdgfrβ-KO obese male (E) and female (F) mice after seven days of cold temperature exposure (n = 4-5 biologically independent mice).

**G, H.** Quantification of adipocyte area from iWAT and gWAT sections from Control and Pdgfrβ-KO obese male (G) and female (H) mice after seven days of CE (n = 3 biologically independent mice).

**I, J.** Quantification of Ucp1 immunostaining of iWAT sections from Control and Pdgfrβ-KO obese male (I) and female (J) mice after seven days of CE (n = 3 biologically independent mice).

**K, L.** Representative images of H&E staining of gWAT sections from Control and Pdgfrβ-KO male (K) and female (L) obese mice maintained cold exposed for seven days.

**M, N.** Relative mRNA levels of denoted thermogenic genes within iWAT depots from cold exposed Control and Pdgfrβ-KO male (M) and female (N) obese mice (n = 4 biologically independent mice).

**O.** Representative flow cytometric histograms of phosphorylated Stat1 within SMA-marked beige APCs from Control and Pdgfrβ-KO male and female obese mice.

Data are means with individual data points ±S.E.M. Statistical significance was determined using paired Student's t test (a and b) and unpaired two-tailed Student's *t* test (C-F, and I-N). Scale bar = 100 μm.

Supplemental Table 1

| Gene | Forward | Reverse |
|---|---|---|
| *IL-13* | *CCTGGCTCTTGCTTGCCTT* | *GGTCTTGTGTGATGTTGCTCA* |
| *IL-33* | TCCAACTCCAAGATTTCCCCG | CATGCAGTAGACATGGCAGAA |
| *Cidea* | *TCTGCAATCCCATGAATGTC* | *CAGTGATTTAAGAGACGCGG* |
| *Dio2* | *ACACTGGAATTGGGAGCATC* | *ATGCTGACCTCAGAAGGGCT* |
| Pdgfrβ | *AGGGGGCGTGATGACTAGG* | *TTCCAGGAGTGATACCAGCTT* |
| *Pgc1α* | *TATGGAGTGACATAGAGTGTGCT* | *CCACTTCAATCCACCCAGAAAG* |
| *SdhD* | *AGACCCGCTTATGTGTCAGC* | *GGAACCAGAGTGGTGGCTTG* |
| *Atp5* | *CGGCCATTTTGTGCCAGTC* | *CATTTTTGGAGACCAGTCCCG* |
| *Ucp1* | *CGACTCAGTCCAAGAGTACTTCTCTT* | *GCCGGCTGAGATCTTGTTTC* |
| *Rn18S* | *GTAACCCGTTGAACCCCATT* | *CCATCCAATCGGTAGTAGCG* |

Primer sequences for qPCR gene expression analysis.
